# The First of Us: *Ophiocordyceps* use a novel scramblase-binding peptide to manipulate zombie ants

**DOI:** 10.1101/2025.09.09.674826

**Authors:** William C. Beckerson, Steffen Werner, Maite Goebbels, João J. Ramalho, Andrew J. M. Swafford, Ilia Soroka, Sinah T. Wingert, Mike Boxem, Suzan Ruijtenberg, Greg J. Stephens, Sander van den Heuvel, Charissa de Bekker

## Abstract

Parasite-adaptive manipulation of behavior is a widespread natural phenomenon. While *Ophiocordyceps* zombie fungi are well-known for hijacking ant behavior to increase their fitness, functionally characterizing the biomolecules involved in behavior manipulation remains difficult in these non-model organisms. To circumvent this problem, we have adopted the powerful genetics toolbox of *Caenorhabditis elegans* to identify molecular targets and neurophysiological effects of candidate *Ophiocordyceps* effectors. With this approach, we discovered a novel cysteine-rich, small secreted fungal peptide that binds to well-conserved calcium-dependent scramblase channels in neurons, particularly those associated with sensory tissues. This binding suppressed nematode motor coordination and dampened ant olfactory systems vital for communication. Our findings are the first to directly connect an *Ophiocordyceps* effector with its extended phenotype, while demonstrating the neuroethological role of Scramblase-1.

## Introduction

Reciprocal coevolutionary pressures experienced between parasites and their hosts can lead to the evolution of intense host specialization and unique infection strategies over time^1^. In some cases, the intimate symbiotic relationship between organisms can link the genome of parasites with phenotypes in its hosts, a phenomenon referred to as an extended phenotype^2–3^. Behavior manipulation is a common example of an extended phenotype found across the tree of life^4^. Using effectors to alter host behavior, parasites can increase their own fitness by augmenting the development and/or transmission of parasitic propagules. Fungal parasites are particularly well known for their behavior manipulation strategies, inspiring pop culture media like *The Last of Us*^5^ and *The Girl with All the Gifts*^6^. The biodiverse species complex *Ophiocordyceps unilateralis*^7–8^ is particularly well characterized in its temporal induction of multiple conspicuous behavioral changes in infected ants. First, the fungus disrupts the social behaviors of its host, reducing their communication with nestmates. Next, the hosts are led to abandon their foraging roles and leave the nest, a behavioral manipulation that helps the parasite avoid detection and destruction by nestmate social immunity responses^9–10^. Infected ants then exhibit summit disease^11^, climbing nearby vegetation to a vantage point that aids in fungal fruiting body development and spore dispersal^12^. Finally, the ant bites down irreversibly to the substrate at this elevated position, ensuring it remains anchored after death when the fungus transitions to its reproductive stage. By stimulating its host to climb to a higher location, the fungus is better positioned to spread its spores more effectively on the wind. While these extended phenotypes may seem uniquely adapted to the life cycle of social ant species, examples of similar behavioral changes (e.g., summit disease) can be found in other parasite-host interactions as well^13–15^. This suggests that many behavioral manipulation strategies have evolved convergently^16^, implying that different parasites may exploit the same core neurobiological pathways in animal hosts. Understanding which pathways are involved and how these pathways are exploited by these parasites is an important step towards understanding unifying neuroethological processes in insects, particularly those with agricultural implications. By exploring behavioral effectors and understanding their impact on conserved insect neurobiology, we open the door to the discovery of new biocontrol agents to help combat destructive pests.

While the high degree of biodiversity and life cycles of behavior-manipulating parasites is well described, molecular studies are often hindered by the lack of bioengineering techniques available in these majoritively non-model organisms. The same is true for many of their hosts and other agricultural pests, which often lose natural behaviors and stress resilience under laboratory conditions^17–18^. While multiomic tools can help identify putative behavioral effectors in some of these parasite-host interactions, characterizing their effects *in vivo* remains difficult. To address this problem, we sought to find a workaround by integrating tools from more genetically tractable model organisms like *Caenorhabditis elegans*. This involved the utilization of a Yeast Two-Hybrid (Y2H) system and several behavioral assays involving *C. elegans*. Using these tools, we were able to reverse engineer the function of a putative zombie-making peptide to inform further studies in the native host. Ultimately, we identified a cysteine-rich, small secreted peptide that binds to calcium-dependent scramblase proteins (SCRM-1 and SCRM-2) in the neurological tissues of nematodes. Cys-rich peptides are well-known contributors to the neurotoxic effects of arachnid venoms^19^, and their capacity to interfere with ion channels^20^ could make them efficient insecticidal candidates^21^.

While scramblase lipid channels have traditionally been implicated in membrane remodeling through phospholipid translocation^22^, more recent findings have challenged this limited scope, proposing other roles when expressed in neurons^23–24^. In both mouse and fly models, scramblases were found to be involved in synaptic transmission, with homologs for scramblases 1 and 2 playing a vital role in neurotransmitter vesicle release and accumulation at neuromuscular junctions^23–24^. Moreover, these scramblases were prominently found in fly antenna^24^, indicating a putative association with invertebrate olfactory sensing. Olfactory disruption during infection could facilitate *Ophiocordyceps*-adaptive abandonment of nestmates and social roles by interfering with the host’s ability to detect odors used by ants to communicate^25^. We therefore hypothesized that this scramblase-binding *Ophiocordyceps* peptide is involved in the nest abandonment behavior of manipulated ants. Through a combination of computational modeling, colocalization imaging, and behavioral experiments, we strengthened our Y2H findings by showing that exposure to this peptide causes olfactory sensing-related behavioral changes in *Camponotus floridanus*. By incorporating the robust molecular toolboxes of model organisms like *C. elegans* into our analysis pipeline, we have demonstrated the power of interdisciplinary research and provide an alternative way to study the neuroethology of behavioral effectors in difficult to work with systems. We have also successfully characterized the first *bona fide Ophiocordyceps* behavioral effector.

## Results

### Identification of a putative Ophiocordyceps effector involved in behavioral manipulation

It has long been hypothesized that *O. camponoti-floridani* manipulates ant behavior using secreted proteins^9^. We therefore compared the secretome of *O. camponoti-floridani*, determined using a rigorous computational analysis^26^ (Data S1), with transcriptomics data to reveal a novel small secreted peptide (Fig. 1A) putatively involved in *Ophiocordyceps* manipulation of ant behavior. This gene (*Ophcf2|06345* in *O. camponoti-floridani*) showed one of the highest degrees of increased expression during manipulated summiting of *C. floridanus* ants when compared to fungal growth in insect medium (log2-fold change = 8.45^9^; Fig. 1B). Its homolog in *O. kimflemingiae* (Ophio5|1675) also shows a similar expression pattern during *Camponotus castaneus* manipulation (log2-fold change = 4.51^27^; Fig. 1B). Moreover, homologs with predicted secretion signals also exist in other behavior manipulating *Ophiocordyceps* species (Fig. 1C), some of which infect other species of insects (e.g., *Ophiocordyceps sinensis* infection of caterpillars^28^). These homologs all share a high degree of sequence conservation in the mature peptide but show variation in the signal peptide regions. Despite this variation, all homologs are predicted to be secreted (SignalP-6.0; Data S2), suggesting that this putative effector might play an important role in general behavior manipulation. Structural predictions of the secreted peptide from *O. camponoti-floridani* with AlphaFold2 (fig. S1) showed configurational parallels with neurotoxic cys-rich peptides^20^, possessing two ShK-like α-helices^29^ and stable antiparallel β-strands that form a disulfide-directed β-hairpin fold^30^. These equidistant cysteines create three tightly connected disulfide bonds characteristic of stable cysteine knots^19–20^ (Fig. 1A). Furthermore, all of the homologs demonstrated strong conservation of these cysteine residues, which accounted for approximately 13% of the mature peptides, with identical spacing maintained across each of the sequences (Fig. 1C). These features provided promising support for further investigation into potential neurological binding partners in the host.

**Fig. 1.**
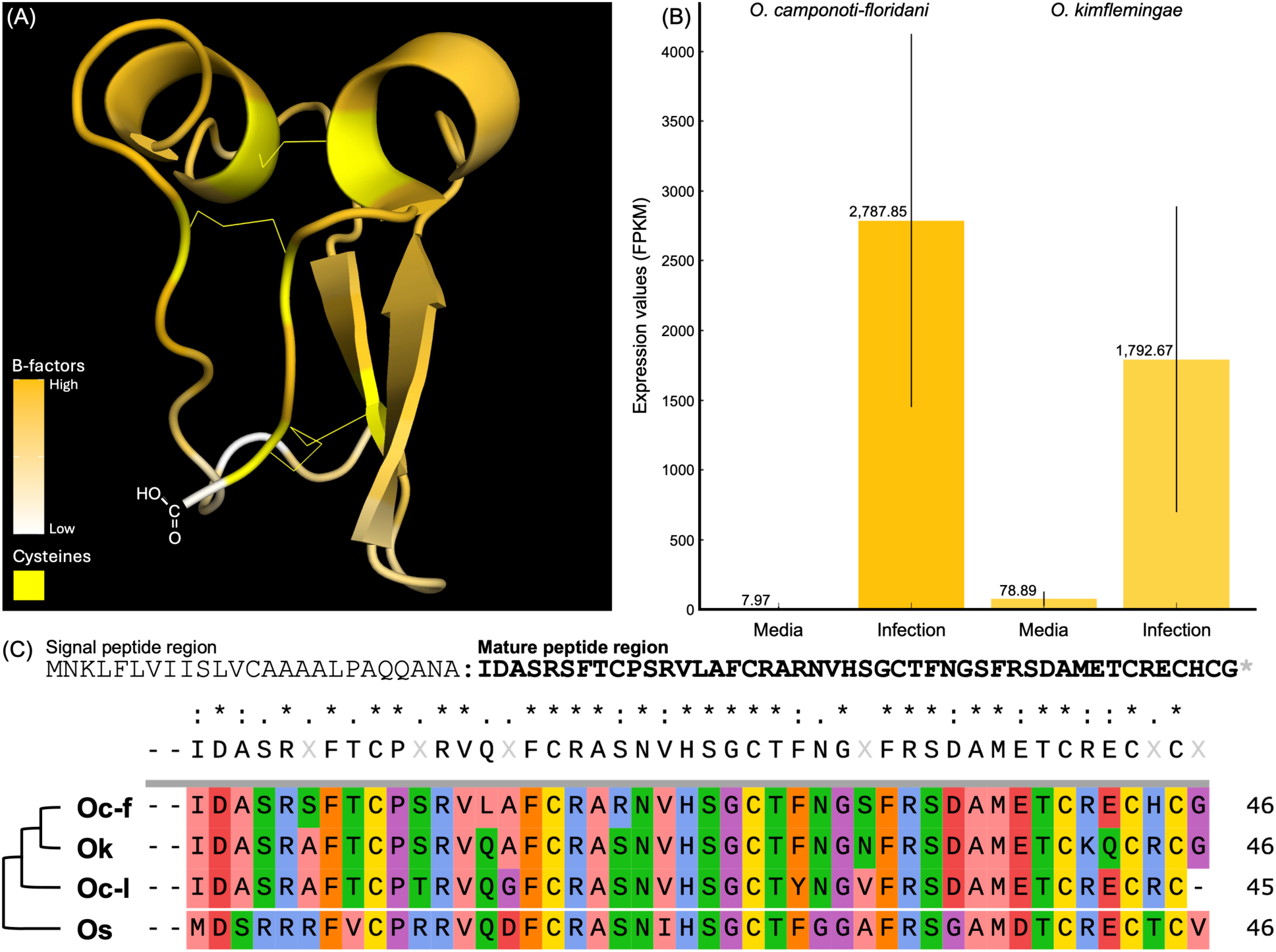
Features of peptide Ophcf2|06345. (A) 3D structural model of Ophcf2|06345 as predicted by AlphaFold2 with high confidence folding shown in orange and low confidence regions shown in white. The position of the cystine residues connected by disulfide bonds are shown in yellow. (B) Expression levels of the Ophcf2|06345 gene in fungi grown in Grace’s insect media compared to infection of carpenter ant hosts. The expression levels for *O. camponoti-floridani* are shown on the left and the expression levels for the homolog in *O. kimflemingae* are shown on the right. (C) Full polypeptide sequence for Ophcf2|06345, top, and the amino acid sequence comparison of the mature peptide regions for homologs found in the *Ophiocordyceps* genus (Oc-f = *O. camponoti-floridani*, Ok = *O. kimflemingae*, Oc-l = *O. camponoti-leonardi*, Os = *O. sinensis*).

### Identification of scramblase lipid channels as a target for binding by the putative Ophiocordyceps effector

To identify candidate binding partners for the Ophcf2|06345 peptide, we performed a series of Y2H assays with cDNA and ORFeome libraries prepared from the *C. elegans* genome. These screens yielded 51 and 44 diploid colonies with robust growth on selective media, respectively (Data S4). Sequencing of prey vectors present in each of these colonies revealed several potential binding partners (Data S4). Of these, an uncharacterized gene predicted to play a role in serine-type endopeptidase inhibitor activity (*Y69H2.3*), genes activated in blocked unfolded protein response (*abu*), and phospholipid scramblases (*scrm*) were found in both libraries with more than two replicates. Given that *abu* genes also commonly appeared in Y2H screens with other *Ophiocordyceps* effector candidates, these interactions were presumably caused by autoactivation and removed from further analysis. We then used BlastP to search the *C. floridanus* proteome (taxid:104421, NCBI) for homologs of the remaining *C. elegans* binding partners. This analysis identified matches for both queries: Y69H2.3 (Camfl2|XM_025412121.1) and SCRM (Camfl2|XM_025407014.1). A closer look at the predicted function revealed that Y69H2.3 homologs had no known function in either worms or ants. In contrast, both of the *C. elegans* SCRM binding partners, SCRM-1 and SCRM-2, aligned with a single ant protein, Phospholipid Scramblase 1 (PS1), which has the same calcium-dependent lipid transporter function. We therefore focused further studies on interactions between *Ophcf2|06345* with SCRM-1 and SCRM-2 (Fig. 2A).

**Fig. 2.**
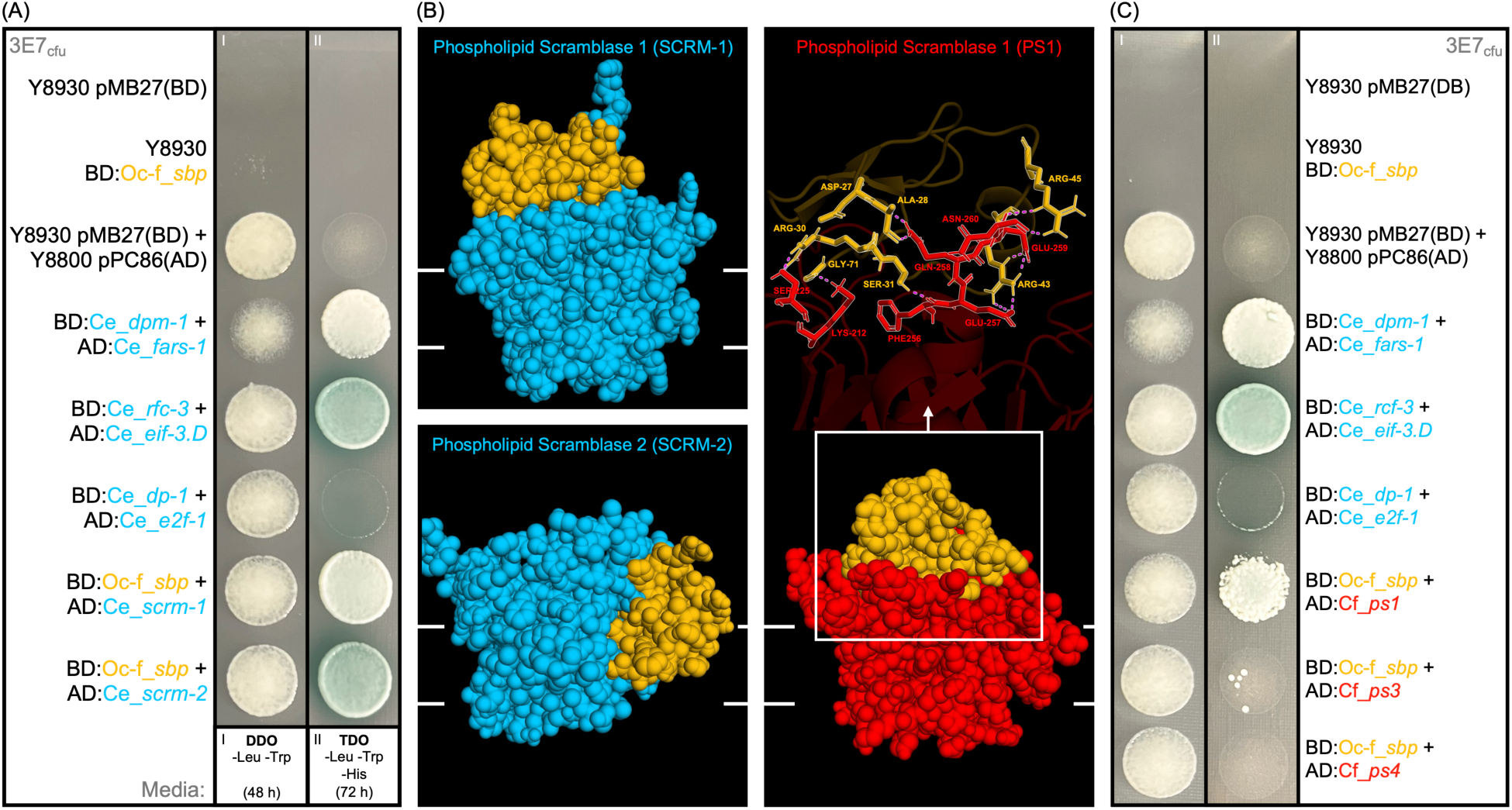
Yeast Two-Hybrid (Y2H) mating assay and computationally predicted binding sites. (A) Mating between yeast strains harboring the coding region for *Ophcf2|06345* protein (SBP) and those harboring the cDNA or ORFeome library from *C. elegans*. The spot assay on the left is plated on double dropout media (DDO) lacking the essential amino acids Leu and Trp to select for successfully mated colonies. The spot assay on the right is plated on triple dropout media (TDO) additionally lacking the amino acid His and with added 3AT to select for robust protein-protein interactions and x-α-Gal to induce a blue colorimetric phenotype. Yeast strains and plasmid names are shown in black text and abbreviated with BD for the binding domain or AD for the activation domain components of the corresponding vectors. *C. elegans* genes are shown in blue, *O. camponoti-floridani* genes in orange, and *C. floridanus* genes in red. Rows 1-2 demonstrate the inability of haploid yeast cells containing the bait vector (pMB27) or the prey vector (pMB27) to grow on either media types. Row 3 demonstrates the ability for diploid mated cells without robust protein-protein interactions to grow on the DDO media, but not the TDO media. Rows 4-6 represent positive controls for the *C. elegans* Y2H libraries, with *dpm-1* x *fars-1* showing weak x-α-gal catabolism, *rfc-3* x *eif-3.D* showing efficient x-α-gal catabolism, and *dp-1* x *ef2-1* showing a very strong rate of x-α-gal catabolism associated with accumulation that affects colony growth. (B) ScanNet predicted binding residues for SBP (orange) with the *C. elegans* scramblases 1 and 2 (blue) and *C. floridanus* scramblase-1 (red). Interacting residues are shown via text overlay alongside their numerical position in each protein. (C) Pairwise mating assay between yeast strains harboring SBP and the *C. floridanus* Phospholipid Scramblases 1, 3, and 4.

In total, nematodes have eight scramblase proteins. This is a comparatively large number compared to other organisms (e.g., five in *Homo sapiens* and *Mus musculus*, and only two in *Drosophila melanogaster*), while *C. floridanus* ants have just three (PS1, PS3, and PS4). We compared the 3D structures of SCRM-1, SCRM-2, and PS1 to determine if the Ophcf2|06345 peptide would likely bind with them as well given that the amino acid similarity of the *C. elegans* scramblases with PS1 is low (35% for SCRM-1 and 36% with SCRM-2, fig. S2). Comparing AlphaFold2 models with Foldseek^31^ and PyMOL v 2.5.8 (Schrödinger, LLC) showed a much higher degree of similarity (75% and 81% TM scores, respectively), particularly in the core region (fig. S2). This structural similarity was also found between PS1 and the two phospholipid scramblases present in the *D. melanogaster* genome (SCRAMB1, 88%, and SCRAMB2, 76%, TM scores; fig. S2). Additional computational modeling of molecular docking between PS1 and the *Ophcf2|06345* peptide, indicates that the peptide binds to the surface of PS1, presumably acting as a cap to interfere with lipid transport (Fig. 2B). Similar results were obtained with SCRM-1; however, SCRM-2 showed a preference for peptide binding from the side (Fig. 2B). While this sort of binding would be possible with proteins in the nucleoplasm during Y2H analyses, it would not occur *in vivo* due to the transmembrane localization of SCRM proteins, making it a weaker candidate. To verify peptide binding to PS1, and to test whether it binds to any of the other ant scramblase proteins, a series of follow-up pairwise Y2H assays was performed between *Ophcf2|06345* and all three phospholipid scramblase proteins from the *C. floridanus* genome. Results showed that *Ophcf2|06345* only binds to PS1 of *C. floridanus*, consistent with our findings from the *C. elegans* libraries (Fig. 2C).

Given that SCRM-2 is expressed in very low quantities across all life stages of nematode development (fig. S3), and that knockout studies in *Drosophila* have demonstrated that repair of *SCRAMB1* alone is enough to rescue wild-type phenotypes in Δ*SCRAMB1;*Δ*SCRAMB2* double mutants^24^, we further narrowed the scope of this study to investigate the relationship between the *Ophcf2|06345* peptide and Scramblase-1 homologs, exclusively. We endogenously tagged SCRM-1 with mKate 2 and expressed a GFP-tagged version of *Ophcf2|06345* in *C. elegans* to demonstrated binding *in vivo.* This was confirmed through a high degree of colocalization in the nervous system of *C. elegans* (Fig. 3). Imaging also showed that SCRM-1 is heavily expressed in the sensory organs (Fig. 3A) and the nerve ring (Fig. 3B; Movies S1-2). Taken together, the co-localization data, alongside Y2H and computational evidence, indicate that this previously undescribed Ophcf2|06345 peptide binds to SCRM-1. We therefore named it the “Scramblase-Binding Peptide” (SBP).

**Fig. 3.**
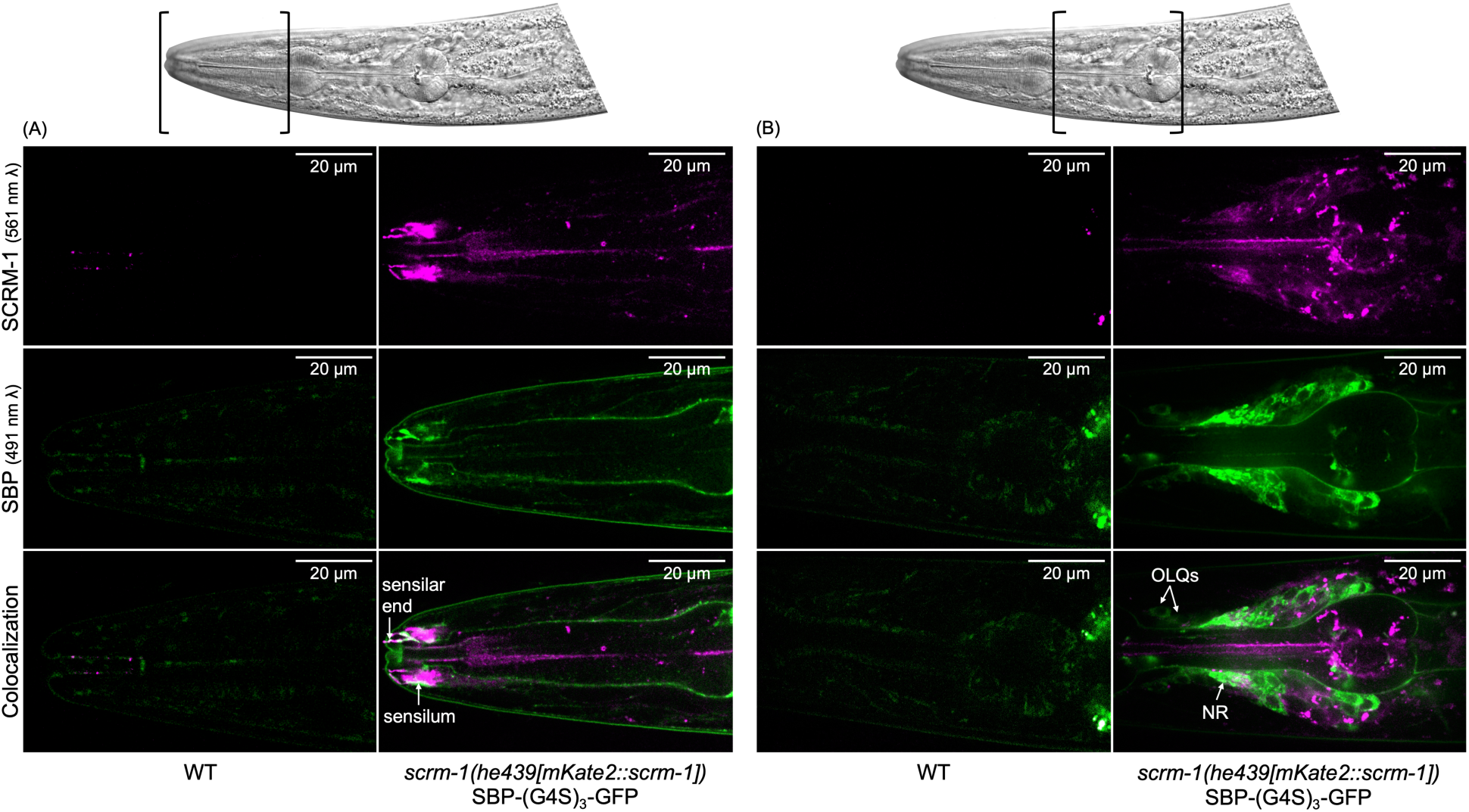
Colocalization of *Ophiocordyceps* Ophcf2|06345 (SBP) and *C. elegans* Scramblase-1. The top images represent spinning disc confocal imagery of the *C. elegans* amphid region, (A), and nerve ring, (B). The top row of fluorescent microscopy images was taken with a 561 nm laser to excite the mKate2 fluorophore linked to SCRM-1, shown in magenta. The middle row of images was taken with a 491 nm laser to excite the GFP fluorophore linked to a codon-optimized variant of SBP, shown in green. The bottom row of images represents a series of composite images with high degrees of colocalization shown in white. The left column in both A and B shows autofluorescence in CGC1 worms while the right columns show fluorescence of the fluorophores in transgenic *scrm-1(he439[mKate2::scrm-1]*) worms expressing pPrab-3_CelOptSBP-(G4S)3-GFP_Tlet-858 (Data S7). Colocalization can be seen to a higher degree in the sensilar tissues, outer labial quadrant neurons (OLQs), and the nerve ring (NR).

### Ophiocordyceps Scramblase-Binding Peptide causes behavioral changes in C. elegans and C. floridanus

To determine if the binding of fungal SBP to SCRM-1 can alter behavioral phenotypes, a series of tests was performed with nematodes and analyzed using our new MATLAB software designed for detecting subtle changes in body position^32^. First, we performed two behavioral experiments involving nematodes exposed to SBP. In these tests, the worms were either 1) fed SBP-expressing Rosetta^TM^ 2 bacteria or 2) genetically modified to express codon-optimized SBP. The results were compared to behavioral tests involving the removal or repression of SCRM-1 using the knockout strain CU2904 (Caenorhabditis Genetics Center) and RNAi^33^, respectively. By comparing the results from all four tests, we aimed to link any behavioral changes caused by SBP, at least in part, with the inhibition of SCRM-1 lipid channel transport. In all four conditions, we observed lower body wave amplitudes during nematode crawling (Movie S3), which are depicted by the ring-shaped probability distribution of the first two modes of a principal component analysis in Fig. 4A and fig. S4. However, despite a consistent lower average diameter in all four groups, only the results from the SBP-exposure tests were statistically significant when compared to their respective controls (two-tailed t-test, p-values 2.32e-3** and 1.91e-8***, respectively; Fig. 4B). These behavioral results, in conjunction with the colocalization assays that show an abundance of SCRM-1 in the nerve ring and sensilar organs of *C. elegans*, suggest that SBP can induce behavioral changes presumably through the inhibition of neuromuscular and/or chemosensory systems. Working from this hypothesis, we conducted further behavioral tests in the native ant hosts, *C. floridanus*.

**Fig. 4.**
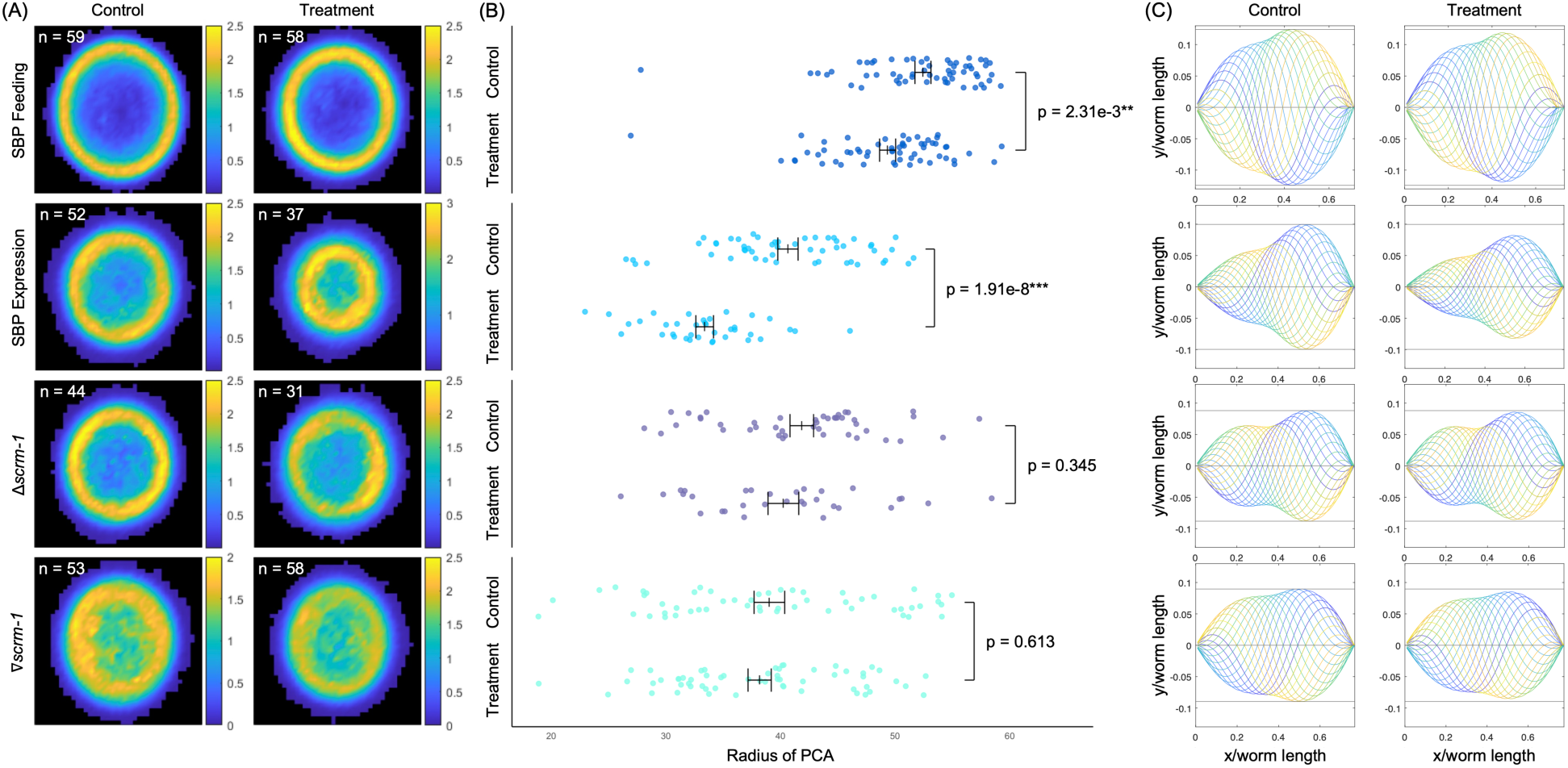
Behavioral analysis of *C. elegans* in response to SBP exposure and SCRM-1 inhibition. The data is organized into rows based on experimental parameters. The top row represents behavioral assays performed with wt nematodes (CGC1) fed SBP-expressing Rosetta^TM^ 2 and controls fed empty vector Rosetta^TM^ 2. The top middle row represents assays performed with transgenic worms self-expressing SBP and transgenic controls expressing GFP, while the bottom middle row presents behavioral data from *scrm-1*-knockout worms (CU2904) and wt control worms. Finally, the bottom row represents KP3948 worms that were fed HT115(DE3) cells expressing dsRNA for *scrm-1* (Vidal RNAi library; Rual et al., 2004) compared to controls were fed cells producing dsRNA for GFP. (A) Principal component analyses (PCA) of the body curvature of crawling *C. elegans* worms. The joint probability distributions of the amplitudes a_1 (x-axis) and a_2 (y-axis) of the first two modes with the largest variances shows a circular structure. This corresponds to travelling waves along the worm body during motion via lateral undulations. Color denotes the relative fraction of data points in ppm. (B) Average radial distance <\sqrt{a_1^2+a_2^2}> in the space of the first two eigenmodes for each worm. Mean and standard deviation are shown for each data set in black. Significance was determined using a two-tailed t-test. (C) Reconstructed worm shapes from the average contributions of only the first two modes of the PCA (fig. S4) illustrate that the smaller radius corresponds to waves with smaller amplitudes, i.e., smaller peak curvatures.

Similar to the *C. elegans* behavioral assays, *C. floridanus* assays were conducted by 1) injecting ants with SBP extracted from the periplasmic space of the same Rosetta^TM^ 2 cells fed to nematodes and 2) repressing PS1 by injecting dsRNA targeting *ps1*. Proper disulfide bond formation for the peptides extracted from the bacterial periplasmic space was verified using a western blot comparing native exacts to those treated with DTT (fig. S5). To record ant behavior, we used a suite of computational tracking software to identify changes in movement and/or chemosensory-related ant behaviors. Compared to controls, both SBP-injected and PS1-repressed ants spent significantly more time near filter pads soaked with citronella oil, a common insect repellant (one-tailed pairwise t-test, p = 2.48e-3** and 0.0121*, respectively; Fig. 5A). These results indicate that the repression of PS1 and exposure to SBP both dampen ant olfactory sensing. This effect was most apparent in the first four minutes of recording, right after the addition of the citronella-soaked pad (fig. S6). Additional pair-wise interaction assays between ants demonstrated that SBP exposure also significantly reduced antennation events compared to controls (two-tailed t-test, p = 0.0285*; Fig. 5B). We also detected a significant reduction in social clustering behavior typically seen between nestmates within the nest, an effect that increased in the SBP-injected group over time (linear mixed-effects model, p < 3.893e-7***; Fig. 5C). Finally, we also observed ants injected with SBP attacking other nestmates during the preparation of behavioral assays (Movie S4), indicating an issue with proper nestmate recognition. Our results, therefore, indicate that *Ophiocordyceps* SBP interferes with host ant chemosensory-related behaviors (e.g., smelling, antennation, and clustering) important for communication and participation in the colony’s social network.

**Fig. 5.**
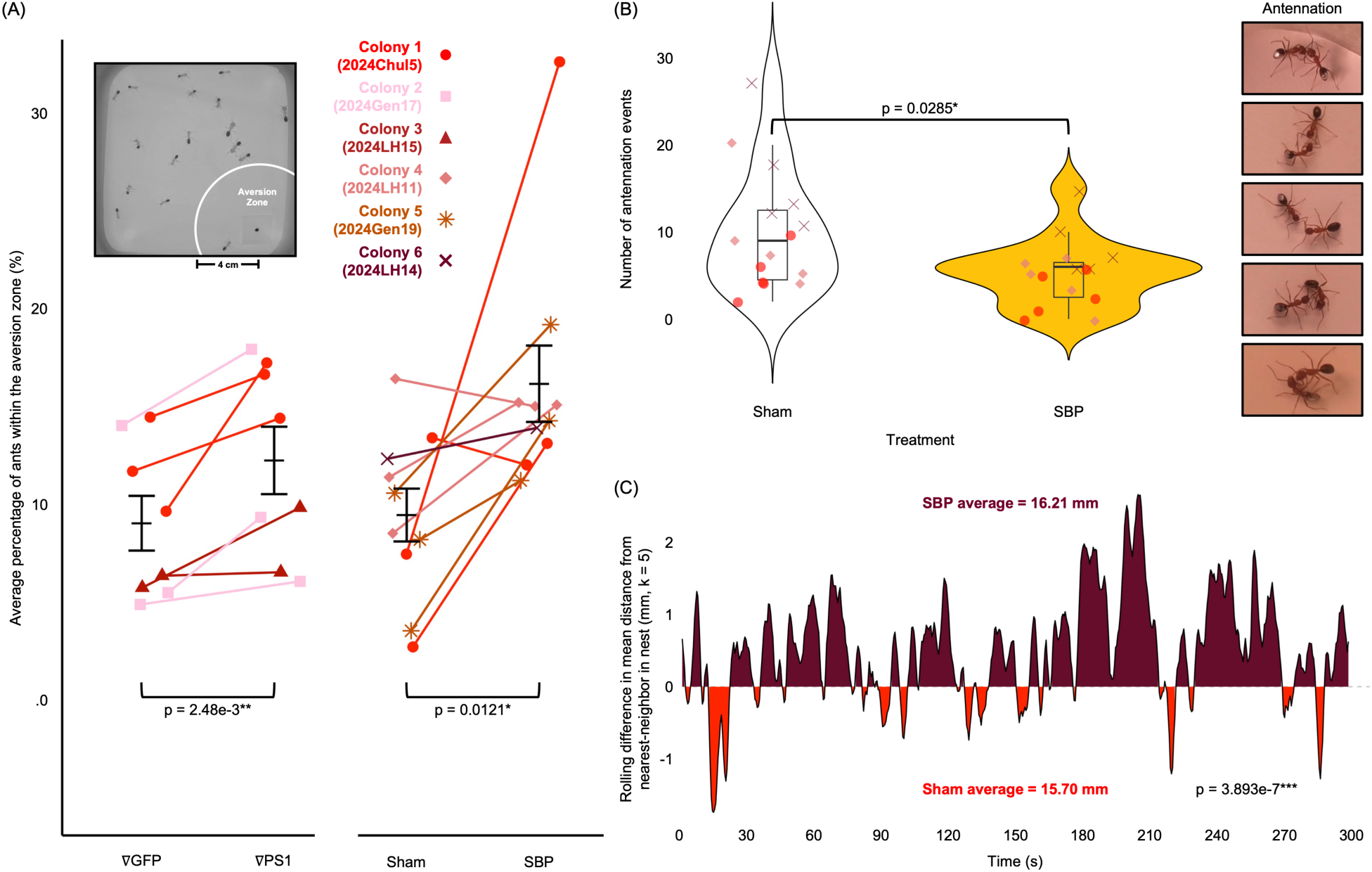
Behavioral analysis of *C. floridanus* in response to SBP exposure and PS-1 repression. (A) The average percentage of ants in the aversion zone (4 cm radius surrounding the citronella pad) calculated as the sum of total time spent by each ant in the zone divided by the total sum time of the recording. Ants injected with dsRNA Phospholipid Scramblase 1 (∇PS1) were compared to those injected with dsRNA for GFP (∇GFP), left. Ants injected with periplasmic protein extracts containing SBP (SBP) produced by Rosetta^TM^2 cells were compared to those injected with periplasmic extracts from expressing an empty plasmid (Sham), right. Statistical significance was determined between the means of each group using a one-tailed paired Student’s t-test. (B) The number of antennation events counted between sham-injected ants and between those injected with SBP, examples of which are shown on the right. Statistical significance between the treatment groups was determined using a two-tailed independent Student’s t-test. (C) A difference area plot for the spatio-temporal distribution of ants in a nest environment after injection with either sham treatment or SBP. Numbers on the positive axis represent a higher degree of separation measured in the SBP-injected ants while numbers on the negative axis represent the same for sham-treated individuals. Measurements were taken every 0.5 sec by calculating the nearest-neighbor distance between each ant averaged across the total number of ants in each frame. The values are shown as a rolling average (k = 5) with time on the x-axis. Statistical significance was determined using a linear mixed-effects model.

## Discussion

While adaptive manipulation of host behavior is a convergently evolved strategy reported across a wide variety of parasite taxa, the molecular interactions that give rise to these extended phenotypes remain largely unknown. In this study, we identified a novel scramblase-binding peptide (SBP) secreted by *Ophiocordyceps*, which appears to dampen carpenter ant olfactory sensing and lessen communication. Carpenter ants rely heavily on antennal sensilla to identify nest mates, communicate with each other, regulate division of labor, and detect sick individuals^34–35^. This fungal SBP may therefore aid parasitic fitness by preventing nestmate recognition^36^ and thwarting recruitment to the nest via pheromones^37^, ultimately disconnecting the host from its social network. Disrupting smell may also prevent the host itself from recognizing changes in its own cuticular hydrocarbon profile during infection^38^, which could prevent self-grooming behaviors or other seemingly altruistic acts of self-sacrifice and kin selection common in eusocial insects due to their high degree of relatedness^39^. Furthermore, without proper functioning olfactory systems, infected insects might instead rely more heavily on visual systems to guide decision making. This could contribute to light-guided summiting behavior that occurs in later stages of infection^11^. By demonstrating that SBP directly induces host behavioral changes in line with important steps of the fungus’s life cycle, we have successfully characterized the first *Ophiocordyceps* behavioral effector and linked it to its extended phenotype in ants.

In addition to the olfactory effects observed in ants, we were able to identify subtle motor effects using *C. elegans* as a surrogate system. Motor coordination dysregulation is another well-defined step of late-stage behavior manipulation in *C. floridanus* infected by *O. camponoti-floridani*^45^. Towards the end of the infection process, ants exhibit “staggers syndrome”, characterized by a loss of coordination and balance, making the ants appear to “stagger” when walking^46^. However, we were unable to detect any movement effects in our ant studies. This is likely due to the difference in sensitivity of the programs used to screen for these effects in each organism. While our analysis of worm movement was performed using our new MATLAB-based program^32^, our preliminary screening for motor problems in ants coopted existing tools that use simple object tracking (i.e., MARGO^40^). Unlike ants, nematodes move with a travelling body wave^41^, which is measurable with wave function mathematics to characterize precise changes in body positions. This allowed us to detect far more subtle differences in nematode body movement compared to our ant analyses. There is however correlative evidence that suggests subtle motor coordination may be present in ants exposed to SBP as well. The abundance of SBPs localization to neuronal tissues is consistent with findings in *Drosophila* that also found expression of Scramblase 1 and 2 homologs at neuromuscular junctions^24^. Furthermore, Scramblase 1 homologs have also been detected in the brains of mice and are implicated in neurotransmission and synaptic vesicle retrieval of the cerebellum^23^. Given that the cerebellum plays a crucial role in muscle coordination, fine motor movements, and linking sensory detection with movement behaviors^42–44^, it makes sense that we detected motor issues in our more sensitive nematode models. The ability to detect such subtle behavioral effects in *C. elegans* underlines the value of using this model organism in the identification of putative behavioral effectors in complex, non-model systems.

With this study, we have taken the largely descriptive research field of neuroparasitology a step further towards characterizing underlying molecular mechanisms by incorporating model systems. This approach provides a new lab-amenable solution for barriers facing the exploration of novel insecticidal compounds from non-model fungal entomopathogens. Furthermore, our findings suggest that scramblase proteins and their role in neurotransmitter release could provide an alternative target for pre/postsynaptic blockade studied in other fields^19–21^, particularly in response to neuropeptides known to bind to various types of ion-dependent channels of the nervous system and cause pain, muscle paralysis, or even death^20,47–48^. The further exploration of neuropeptides like SBP and their effect on odor-mediated communication in social insects presents a new avenue for the discovery of insecticidal compounds and alternative biocontrol solutions that may spare natural fauna. This work therefore provides a path for screening proteomes for novel effectors and testing their effects on invertebrate nervous systems, particularly those without previously defined function.

## General

We would like to thank The Lili’s Proto Lab at Utrecht University for assistance in printing the replica plating apparatus used for the Y2H analyses, Savvas Tzavellas for assistance with setting up and running The AlphaFold Colab notebook (GitHub: AlphaFold.ipynb), Ruben Schmidt for training with the DIC microscope, and Robin Jonkergouw and Emmeline Roosmalen for collecting ants later used as part of this research. The knockout strain CU2904 was provided by the Caenorhabditis Genetics Center (CGC) funded by the NIH Office of Research Infrastructure Programs (P40 OD010440).

## Author contributions

1. Conceptualization: WCB, CdB
2. Methodology: WCB, SW, MG, AJMS, STW, MB, SR, GJS, SvdH, CdB
3. Software: SW, AJMS, GJS
4. Validation: WCB, SW, MG, JJR, AJMS
5. Formal Analysis: WCB, SW, MG, JJR, AJMS, IS, CdB
6. Investigation: WCB, SW, MG, JJR, AJMS, IS
7. Resources: WCB, SW, JJR, MB, SR, SvdH, CdB
8. Data Curation: WCB
9. Writing – Original Draft: WCB
10. Writing – Review & Editing: WCB, SW, MG, JJR, AJMS, IL, STW, MB, SR, GJS, CdB
11. Visualization: WCB, SW, MG, JJR, AJMS, IS, GJS
12. Supervision: WCB, SvdH, CdB
13. Project Administration: WCB
14. Funding Acquisition: WCB, CdB

## Diversity, equity, ethics, and inclusion

All experiments performed as part of this study were conducted following ethical guidelines for animal research. This included upholding the 3R standards; 1) *Replacing* the use of living invertebrates with alternatives like computer models and cell cultures where possible, 2) *Reducing* the number of invertebrates used in experiments to the smallest sample size necessary to achieve statistically significant results, and 3) *Refinement* of protocols to minimize distress or harm caused to invertebrates both during experimentation and during their prior housing.

To make the resulting data and figures more accessible for all audiences, the authors also used a custom colorblind-friendly palette (Table S1) based on prior work published by Wong et al.^49^ A separate color scheme was used for the coexpression analysis based on Fiji options (Table S2). Furthermore, to ensure accurate and ethical demonstration of results, figures involving the use of color gradients used a color scheme with perceptual uniformity following guidelines published by Crameri et al.^50^

## Data and materials availability

Files for 3D printing the replica plating apparatus used in this study can be found at https://github.com/WCBeckerson/3D-Print-Replica-Plating-Apparatus. All 512 nematode and 88 ant video files analyzed in this study are deposited at Zenodo under the DOIs <u>10.5281/zenodo.16760960</u> and <u>10.5281/zenodo.16760416</u>, respectively.

## Tables

N/A

## Supplementary Materials for

### The PDF file includes

Materials and Methods

Supplementary Text

Figs. S1 to S8

Tables S1 to S5

References

### Other Supplementary Materials for this manuscript include the following

Movies S1 to S4

Data S1 to S12

## Materials and Methods

### Computational predictions of Ophiocordyceps camponoti-floridani effectors

#### Bioinformatic screening for secreted effectors

The *O. camponoti-floridani* secretome predicted by Will et al.^9^ using SignalP v4.1^51^ was further analyzed using SignalP v6.0^52^, TMHMM 2.0^53^ Phobius^54^, Prosite^55^, PredGPI^56^, NucPred^57^, and TargetP v2.0^58^ to create a more conservative secretome. This approach reduced the number of predicted secreted proteins from 801 to 586 (Data S1). The remaining 586 candidates were further screened for upregulation in the heads of *Camponotus floridanus* ants during manipulated summiting behavior compared to fungal growth in Grace’s insect media (Sigma) using the RNAseq data from Will et al.^9^. While several genes were found to be highly upregulated during infection, the expression levels of *Ophcf2|06234* (Scramblase-1-Binding Peptide “SBP”) were particularly noteworthy with one of the highest log2 fold changes in the secretome. We used BLASTp analysis (online BLAST+ 2.15.0 tool^59^) to align this peptide against the National Center for Biotechnology Information (NCBI) library to investigate the presence of homologs in other members of the Ophiocordycipitaceae family (Fig. 1A).

#### AlphaFold2 modeling of SBP

The full-length amino acid sequence for SBP was used to predict three-dimensional folding using the default parameters of AlphaFold2 v2.3.1 via ColabFold v1.5.3^60^. The resulting. PDB files were visualized using PyMOL v 2.5.8 (Schrödinger, LLC). The SBP model showed a high level of folding confidence for the structural region of the peptide, but a low confidence for the secretion signal region (fig. S2). Because this region is cleaved off during transport of the protein across the fungal membrane, it was removed for subsequent structural analyses (Fig. 1A).

### Yeast Two-Hybrid mating assays

#### C. elegans cDNA and ORFeome library screening with SBP

The mature peptide region of *SBP* was amplified from cDNA samples of *O. camponoti-floridani*-infected *C. floridanus* using the primers pMB27_Ophcf2|06345 and pMB27_Ophcf2|06345-Rv (Data S7), synthesized by Biolegio (Nijmegen, NL), using Q5® High-Fidelity DNA Polymerase (New England Biolabs). The primers included 20 bp of additional sequence at the 5’ ends complementary to regions of the Y2H bait vector pMB27 (pPC97+4xGly+AscI-NotI) just up and downstream of the restriction sites AscI and NotI, respectively (Data S8). Gibson Overlap PCR using the NEBuilder® HiFi DNA Assembly Cloning Kit (New England Biolabs) was performed to assemble the amplicon with AscI– and NotI-digested pMB27, placing the insert downstream and in-frame of the encoded GAL4 DNA binding domain.

The resulting plasmid pMB27_Ophcf2|06345 (Data S8) was transformed into the *S. cerevisiae* strain Y8930^61^ (*MATα trp1−901 leu2−3,112 ura3−52 his3−200 gal4Δ gal80Δ cyh2R GAL1::HIS3@LYS2 GAL2::ADE2 GAL7::LacZ@met2*) using the Frozen-EZ Yeast Transformation II Kit (Zymo Research). Putative transformants were selected on single dropout media (SDO) lacking leucine (SDO_-Leu_) before verification through yeast colony PCR with the primers Y2H Bait Screen Fw & Rv (Data S7). All dropout media used in this study also contained 100 μg/mL ampicillin to help prevent contamination by bacteria. Verified Y8930 transformants were mated against Y8800 cells (*MATa trp1−901 leu2−3,112 ura3−52 his3− 200 gal4Δ gal80Δ cyh2R GAL1::HIS3@LYS2 GAL2::ADE2 GAL7::LacZ@met2*) housing either the cDNA^62^ or ORFeome^63^ prey vector libraries for *C. elegans* cloned into the multiple cloning site (MCS) of pPC86 (Asc-Not + CyH2) (Data S8). Y2H mating assays were performed based on the protocol published by Li et al.^64^, which includes multiple screening options to help rule out weak interactions and false positives. Diploids colonies were considered to contain strong protein-protein interactions when the following conditions were observed: 1) Robust growth on media lacking adenosine, 2) Robust growth on media lacking histidine, 3) Robust growth on media lacking histidine and including the competitive binding 3AT, and 4) a lack of growth on media containing cycloheximide^64^. Notably, blue/white screening is not as indicative of strong protein-protein interaction for these strains as it is in other Y2H kits (e.g., the Matchmaker^TM^ Yeast Two-Hybrid system^65^).

To screen for protein-protein interactions, mated colonies were first plated on triple dropout (TDO) media lacking the amino acids leucine, tryptophan, and histidine (TDO _-Leu –Trp –His_). These plates were incubated at 30°C for 72 h to allow for sufficient growth under stressful conditions. To calculate mating efficiency, a 10,000X diluted sample of the mating suspension was also plated onto double dropout (DDO) media only lacking leucine and tryptophan (DDO _-Leu –Trp_), which were incubated for 48 h before colony counting. Colonies that grew on the TDO _-Leu –Trp –His_ plates were replica plated onto the following four additional media types to screen for robust protein-protein interactions; 1) TDO _-Leu –Trp –His_ containing 2 mM 3-AT, 2) TDO _-Leu –Trp –Ade_, 3) DDO _-Leu –His_ containing 1 μg/mL cyclohexamide, and 4) DDO _-Leu –Trp_. Replica plating was performed using cotton velveteen cloth and a custom 3D-printed 94 mm diameter replica plating apparatus (https://github.com/WCBeckerson/3D-Print-Replica-Plating-Apparatus).

Colonies that demonstrated robust growth on the first plate, white color on the second plate, no growth on the third plate, and growth on the fourth plate were considered robust interactions and selected for retrieval of the prey vector. The coding region in each of these prey vectors was amplified via yeast colony PCR (DreamTaq, ThermoFisher Scientific) and sent for sequencing (Macrogen) using the universal primers Y2H_Prey_Screen-Fw and Y2H_Prey_Screen-Rv (Data S7) to identify the protein-protein binding partners of SBP. Because the length of the encoded gene in each pPC86 (Asc-Not + CyH2) plasmid (Data S8) was unknown, an extension period of 3 min was used in all PCR reactions. Any binding partner sequence that was not identified more than two times, in both libraries, or which was identified in other Y2H assays for unrelated bait genes, was considered a false positive due to autoactivation or other phenomena and removed from further consideration. These screening criteria resulted in two unique and robust potential protein family binding partners for SBP; Y69H2.3 and the Scramblases SCRM-1 and SCRM-2 (Data S4).

#### Pairwise assay between SBP and C. floridanus scramblase homologs

BLASTp analysis of the *C. floridanus* genome for homologs of the *C. elegans* SCRM-1 and SCRM-2 interaction partners was performed using NCBI. Results showed a shared match for both nematode scramblases with the ant Phospholipid Scramblase 1 (PS1) (Camfl2|XM_025407014.1). To verify if protein-protein interaction also occurred between the SBP and *C. floridanus* PS1 homolog, we amplified the coding sequence for PS1 from cDNA of *C. floridanus* using the primers PS1 Fw and PS1 Rv (Data S7) for cloning into the pPC86 (Asc-Not + CyH2) prey vector (Data S8). The resulting plasmid was named pPC86_PS1 (Data S8). To investigate binding specificity, we also created prey vectors for the other two phospholipid scramblase genes present in the *C. floridanus* genome (PS3 and PS4) using the primers PS3 Fw and PS3 Rv, and PS4 Fw and PS4 Rv to generate the vectors pPC86_PS3 and pPC86_PS4, respectively (Data S8). All three constructs were transformed into Y8800 cells and mated against the same Y8930 strain containing the SBP prey vector used for Y2H against the *C. elegans* libraries (Fig. 2C).

### Computational analysis of Scramblase homologs

#### Bioinformatic comparison of Scramblase homologs

While BLASTp alignments resulted in comparatively low sequence similarities between ant PS1 and the nematode SCRM-1 and SCRM-2 proteins, comparisons of predicted 3D protein models showed a much higher degree of similarity (fig. S3). This was determined by comparing the predicted 3D structures of scramblase homologs of *C. floridanus* (PS1; AF-E2ATE7), *C. elegans* (SCRM-1; AF-O45799 and SCRM-2; AF-G5EEQ3), and *D. melanogaster* (SCRAMB1; AF-Q9VT88 and SCRAMB2; AF-Q9VZW1) using the PyMOL v 2.5.7 (Schrödinger, LLC) align command for seq alignment with structural superposition.

#### SBP-Scramblase binding residue predictions

ScanNet^66^ was used to predict protein binding regions of the 3D models for SBP and scramblase binding partners confirmed by Y2H assays (SCRM-1, SCRM-2, and PS1). Surface residues predicted to have binding propensity scores exceeding 50% were selected as potential interaction sites and used to define active regions for protein-peptide docking. Molecular docking simulations were performed using the standard protocol in HADDOCK v2.4 with resulting clusters scored and ranked by HADDOCK^67^. The three top-ranked models were further analyzed for interface contacts (Fig 2B).

### Co-localization of SBP with SCRM-1 in C. elegans

#### CRISPR-Cas9-mediated tagging of SCRM-1 with mKate2

To visualize the internal localization of SCRM-1 in nematodes, transgenic worms were created using CRISPR-Cas9-mediated insertion of the mKate2 cassette at the N-terminus of the SCRM-1 coding region. The N-terminus was chosen to ensure that all four transcript variants of *scrm-1* were tagged with the fluorescent marker. This approach is also in line with methods used by Acharya et al. to identify SCRAMB1 localization in fruit flies^24^. The *scrm-1* gene was edited at the +4 position using 0.5 μL of 10 μg/μL *S. pyogenes* Cas9-3NLS protein combined *ex vivo* with 5 μL of universal 0.4 μg/μL tracrRNA (Integrated DNA Technologies) and 2.8 μL of the 0.4 μg/μL crRNA SCRM1 target seq (Data S7). The mixture was then incubated at 37°C for 15 minutes before a repair template was added to a final concentration of 25 ng/μL. The repair template was designed by amplifying the mKate2 gene from pMS01 Pmcm-4_cdk-2 sensor_mKate2_tbb-2UTR_ttTi5605 (Data S8) with Q5^®^ High-Fidelity DNA Polymerase (New England Biolabs). To facilitate homologous recombination *in vivo*, the primers used to amplify mKate2, SCRM1-mKate2 Fw and SCRM1-mKate2 Rv, were designed with 35 bp of 5’ complementary sequence to either side of the CRISPR-Cas9 incision point. Following amplification via PCR, the amplicon was denatured and reannealed using the following stepwise thermocycler program before its addition to the Cas9 complex: 95°C – 2:00 min, 85°C – 10 sec, 75°C – 10 sec, 65°C – 10 sec, 55°C – 30 sec, 45°C – 30 sec, 35°C – 10 sec, 25°C – 10 sec, 4°C – hold. The resulting Cas9 complex and repair template were then centrifuged together at max speed for 2 min and the supernatant was moved to a new tube. This was done to pellet any particulate matter in the tube before proceeding with transformation via microinjection of CGC1 worms.

Putatively transformed nematodes were isolated individually on NGM plates seeded with OP50 and allowed to recover overnight at 20°C in a dark incubator. F1 progeny from each putative transformant were lysed to extract genomic material to screen for the presence of mKate2 using PCR and the primers SCRM1-mKate2 Seq Fw and SCRM1-mKate2 Seq Rv (Data S7). Worms that showed bands around 1,200 bp and 350 bp, indicative of heterozygous insertion, were selected and allowed to reproduce for a further three generations. The F4 progenies were then re-screened with the same PCR approach to look for transformants with a single band around 1,200 bp, indicating homozygous fixation of the mKate2 gene. These homozygous transformants were sent for sequencing (Macrogen) to identify worms with proper in-frame insertion of the *mKate2* sequence at the 5’ end of the *scrm-1* gene. A single successful transformant was selected and named *scrm-1*(he439[*mKate2::scrm-1*]) (strain name: SV2604). Progeny of this transformant were used for further transformation with the GFP-tagged SBP expression vector for colocalization assays (Fig. 3).

#### Plasmid creation for expression of GFP-tagged SBP in C. elegans

To build an expression vector for GFP-tagged SBP in *C. elegans*, the mature peptide sequence from *O. camponoti-floridani* was codon optimized using WormBuilder Transgene Adaptation Beta (WormBuilder; Data S9). The endogenous signal peptide region was replaced with the nematode secretion signal *spp-1*^68^ (Data S9). The resulting sequence was checked against the IDT gBlocks^TM^ Gene Fragments Entry Tool (Integrated DNA Technologies) and ordered as a single oligonucleotide from GeneArt Strings (Thermo Fisher Scientific). This “Codon Optimized SBP” fragment (Data S7) was bridged with the *rab-3* promoter for post-differentiation pan-neuronal expression to mimic the measured expression of *SBP* in ant heads during infection and manipulation by *Ophiocordyceps* fungi. The *let-858* terminator was added to the 3’ end of the sequence via Gibson Overlap PCR with the NEBuilder^®^ HiFi DNA Assembly Kit (New England Biolabs). Both the promoter and terminator fragments were amplified using Q5^®^ High-Fidelity DNA Polymerase (New England Biolabs) and the primer pairs Gib_rab-3 Fw/Gib_rab-3 Rv and Gib_let-858 Fw/Gib_let-858 Rv, respectively (Data S7). The resulting plasmid was further augmented with a GFP fluorophore added to the 3’ end of the coding region. The sequence for GFP was amplified from the vector pMyo2_GFP (Data S8) with the primers (G4S)3-CelOptOphcf2|06345 RV and (G4S)3-CelGFP Fw (Data S7). These primers contained an additional sequence that introduced a (G4S)_3_ linker bridging the GFP sequence with the codon-optimized SBP sequence. The resulting plasmid was named pPrab-3_CelOptSBP-(G4S)3-GFP_Tlet-858 (Data S8) and injected into transgenic *scrm-1*(he439[*mKate2::scrm-1*]) worms for co-localization assays using fluorescent microscopy (Fig. 3).

#### Microinjection of C. elegans

To prepare young adult worms for microinjection, L4 worms were moved to an OP50-seeded NGM plate the day before injection and incubated overnight at 20°C. Injection pads were prepared by flattening a drop of 2% agarose (Fisher bioreagents) between two coverslips. Microcapillary Borosilicate glass needles (World Precision Instruments) were prepared by pulling the glass microcapillaries in a Model P-1000 Flaming/Brown micropipette puller (Sutter Instrument) set to the following parameters: 1 Line, Heat – ramp to 480, Pull – 50, Vel – 60, Delay – 90, Pressure – 200, 2.5 x 2.5 mm Box, 1 x 0.75 mm. Needle tips were opened by dragging them gently across the length of another glass capillary tube under halocarbon oil in a lateral, perpendicular motion.

To inject, young adult worms were transferred to a droplet of ∼5 μL halocarbon oil 700 (Sigma) on a 2% agarose pad using a size 15 series 170 MICRO-NOVA synthetic fiber brush (da Vinci Brush Maker). The brush was also used to straighten and embed them into the agarose pad. Once positioned, we proceeded with microinjection using a high-resolution Observer.A1 Axio confocal microscope (Zeiss) equipped with a TransferMan^®^ 4r remote-controlled robotic injecting arm (Eppendorf) and a FemtoJet 4x microinjector (Eppendorf). Injections were performed in the distal regions of both gonad arms with the following microinjector settings: injection pressure (pi) – 28.03 PSI, injection time (ti) – 0.72 s, and compensation pressure (pc) – 0.40 PSI. These settings were slightly adjusted for each series of injections depending on the degree of breakage in the needle tip.

Preliminary analysis of pPrab-3_CelOptSBP-(G4S)3-GFP_Tlet-858 transformations indicated that the GFP-tagged SBP did not congregate anywhere in high enough abundance to be used as a screening marker for transformation. Therefore, injection of worms with this plasmid were performed with the addition of pRF4 helper plasmid (Data S8) expressing *rol-6*(su1006) that produces a dominant roller phenotype as a co-injection marker. After injection, each worm was placed onto its own individual NGM plate seeded with OP50 and freed from the halocarbon oil using a 5 μL droplet of M9 buffer (22 mM KH2PO4, 42 mM Na2HPO4, 86 mM NaCl, 1 mM MgSO4) before being incubated overnight at 20°C in the dark to recover and reproduce. Putatively transformed progeny in the F1 generation were screened for selectable phenotypes (GFP in the case of pPrab-3_CelOptSBP-T2A-GFP_Tlet-858 or the roller phenotype in the case of pPrab-3_CelOptSBP-(G4S)3-GFP_Tlet-858) over the next couple days as they matured. All transformants were further verified with PCR using DNA extract from single worm lysates.

#### Fluorescent microscopy

Prior to imaging, all *C. elegans* strains were cultured following standard conditions^69^ using hermaphrodites grown at 20°C on nematode growth medium (NGM) agar plates. Imaging was performed on the following strains: wt (CGC1), scrm-1(he439[mKate2::scrm-1]) I (SV2604), heEx618[SBP-GFP, pRF4]; scrm-1(he439) line1 (SV2606), and heEx619[SBP-GFP, pRF4]; scrm-1(he439) line2 (SV2607). Larvae and adults were paralyzed using 20 mM tetramisole solution in M9 buffer and mounted on glass slides coated with pads of 5% agarose in M9 buffer. Confocal spinning disk microscopy was performed with a Nikon Eclipse Ti2-E microscope equipped with a Yokogawa CSU-X1-M1 spinning disk, an Omicron LightHUB 6 laser box (GFP excitation 488 (4.48 mW), mKate2 excitation 561 (3.18 mW)), Nikon CFI Plan Apo lambda 60x/1.4 NA or 100x/1.45 NA oil immersion objectives, and a PrimeBSI sCMOS camera for signal detection. During acquisition, the microscope was controlled using MetaMorph software (v7.10; Molecular Devices) and settings were kept constant in animals of each strain. Epifluorescence microscopy was performed using an Axioplan2 upright microscope (Zeiss), equipped with a DIC polarizer, an Axiocam MRm CCD monochrome camera, and a Plan-Apochromat 63×/1.4 Oil DIC M27 objective.

### Behavioral tests in C. elegans

#### Age standardization of worms

Age synchronization was implemented by picking 15 adult worms onto fresh NGM media seeded with OP50 5 days prior to recording and letting them grow until the next generation reached their egg-laying stage (78 h). The plates were then washed with M9 solution and pelleted via centrifugation for 10 sec at 360 x g. Once the worms were at the bottom of the centrifugation tubes, the supernatant was removed and replaced with 2 mL of 0.25 M NaOH 1% bleach solution. The bleach/worm suspension was vortexed for 60 sec before repeating centrifugation to re-pellet dead worms. The bleach supernatant was then replaced with 2 mL of fresh bleach solution and vortexed again until approximately 2/3 of the adult worms had dissolved, releasing their eggs into suspension. The remaining eggs and worm fragments were pelleted and washed four times with 5 mL of M9 solution to remove the bleach before resuspension in 2 mL of M9. The resulting egg suspension was transferred to a Sterile CELLSTAR^®^ PS, 60 x 15 mm, vented culture dish (Greiner) for incubation at 20°C overnight (15 h). The next morning, the culture dishes were gently agitated to dislodge L1 paused worms and filtered through a 20 mM Hydrophilic, Nonsterile Nylon Net (Sigma-Aldrich) to remove any remaining unhatched eggs. The flowthrough was collected in a 15 mL Falcon tube and subsequently centrifuged for 10 sec at 360 x g to pellet the L1 worms before removing most of the supernatant, leaving 100 μL. The remaining 100 μL of worm suspension was then transferred to a fresh NGM plate seeded with OP50 and returned to the 20°C incubator for 30 h to allow the now synchronized L1 worms to develop into their L4 stage for recording.

#### Heterologous production and feeding of SBP to C. elegans

The *O. camponoti-floridani* SBP was heterologously produced in Rosetta^TM^ 2 *E. coli* cells. This was done using the pET-22b vector containing an IPTG-inducible T7 promoter (Data S8). The native sequence for SBP without its signal peptide region was amplified from cDNA of *O. camponoti-floridani*-infected *C. floridanus* using Q5® High-Fidelity DNA Polymerase (New England Biolabs) PCR and the primers SBP_NoSig Fw and SBP_NoSig RV (Data S7). The resulting amplicon was cloned into the MCS of pET-22b digested with NcoI and NotI (New England Biolabs) using the NEBuilder^®^ HiFi DNA Assembly Kit (New England Biolabs). This placed the gene in frame with a *pelB* signal sequence for transport of the translated SBP to the periplasmic space. The periplasmic space provides an oxidizing environment that is important for proper disulfide bond formation and folding of eukaryotic proteins. Furthermore, a 6xHis tag was added to the C-terminus of the protein for verification of expression via Western Blot. Cloning was facilitated by the NEBuilder^®^ HiFi DNA Assembly Kit (New England Biolabs). The resulting pET-22B_SBP-6xHis vector (Data S8) was then transformed into Rosetta^TM^ 2 (DE3) Competent Cells (Novagen) using conventional CaCl_2_-treated competent cells and subsequent heat shock transformation. Putative transformants were identified via colony PCR and verified using whole plasmid nanopore sequencing (Plasmidsaurus).

To express SBP, cells were grown in a 30 mL tube (Dispolab) containing 25 mL of LB media with 100 μg/mL ampicillin and 25 μg/mL chloramphenicol to carrying capacity by shaking at 200 rpm in a 37°C incubator overnight for ∼16 h. In the morning, IPTG (dioxane-free, Thermo-Scientific) was added to a final concentration of 1 mM and the temperature was reduced to 28°C.

After 1 h of IPTG induction, cells were resuspended to an OD_600_ = 5.0 in fresh media and 250 μL of the cell suspension was spotted onto NGM plates containing 100 μg/mL ampicillin, 25 μg/mL chloramphenicol, and 1 mM IPTG (dioxane-free, Thermo-Scientific). These plates were stored at room temperature in the dark for 48 h to dry, and the bacteria were allowed to grow into a thin lawn before adding worms. Age-synchronized CGC1 worms in L1 pause were then transferred to these plates and incubated in the dark at 20°C for 30 h before behavioral observations. The behavior of these worms was compared to worms fed Rosetta^TM^ 2 bacteria transformed with an empty pET-22b vector (Data S8), also grown on NGM plates containing 100 μg/mL ampicillin, 25 μg/mL chloramphenicol, and 1 mM IPTG (dioxane-free, Thermo-Scientific).

#### Confirmation of disulfide bond formation in Rosetta^TM^ 2-produced SBP

To test if the disulfide bonds were properly forming in SBP produced by the Rosetta^TM^ 2 expression strains, we used an assay based on work performed on *Schizophyllum commune* proteins^70^. Protein samples were prepared with 4X Laemmli sample buffer, with or without 50 mM DTT and boiled for 10 min. These samples were resolved by electrophoresis on 12% SDS-PAGE gels. Following electrophoresis, proteins were transferred onto a nitrocellulose membrane for immunoblotting. Membranes were probed with a mouse monoclonal 6x-His tag antibody (clone MA-21315, Invitrogen), followed by incubation with a horseradish peroxidase (HRP)-conjugated goat anti-mouse secondary antibody (A16072, Invitrogen). Chemiluminescent detection was performed using the Clarity™ Western ECL Substrate (Bio-Rad) according to manufacturer’s instructions. Blots were visualized using a Bio-Rad Gel Doc Imaging system (fig. S5).

#### Heterologous expression of O. camponoti-floridani SBP in C. elegans

Expression of SBP in nematodes for use in behavioral tests was facilitated by an alternative version of the pPrab-3_CelOptSBP-(G4S)3-GFP_Tlet-858 vector created by replacing the (G4S)_3_ linker with a T2A linker. This plasmid was named pPrab-3_CelOptSBP-T2A-GFP_Tlet-858 (Data S8). This T2A linker induces ribosomal skipping, releasing the GFP and allowing it to accumulate in the cell whilst the SBP is secreted and free of allosteric effects from the comparatively much larger fluorescent tag (SBP 7.64 kDa; GFP fluorophore 27 kDa). Cloning of this plasmid was done using the same method used for the generation of pPrab-3_CelOptSBP-(G4S)3-GFP_Tlet-858 but with the primers T2A-CelGFP Fw and T2A-CelOptOphcf2|06345 RV (Data S7). pPrab-3_CelOptSBP-T2A-GFP_Tlet-858 was subsequently transformed into CGC1 worms via microinjection and preliminary screening of transformants indicated that the concentration of GFP maintained within the cells was high enough for robust selection of transformants (fig. S7). These SBP-expressing worms were compared with transgenic CGC1 worms transformed with a truncated version of the plasmid, pPrab-3_GFP_Tlet-858, expressing and secreting the GFP region without the SBP (Data S8). Both transformants were grown on OP50-seeded NGM plates prior to behavioral tests.

#### C. elegans SCRM-1 knockout

The nematode strain CU2904 with genotype scrm-1(tm698) I. was ordered from the Caenorhabditis Genetics Center (CGC) for use in behavioral tests. This strain was passaged for four consecutive generations under non-stressful growth conditions (OP50-seeded NGM plates, 20°C, no longer than 3 days to prevent overcrowding) before conducting behavioral experiments to help eliminate any transgenerational effects of stress^71^. These worms were compared to CGC1 worms also grown on OP50-seeded NGM plates.

#### dsRNA-induced down regulation of SCRM-1 in C. elegans

To test the behaviors of CGC1 worms with functional but down regulated SCRM-1 proteins, worms were fed HT115 (DE3) bacteria producing dsRNA for *C. elegans scrm-1* from the Vidal library^33^. Because RNAi is less effective in the neurological tissue of wild type *C. elegans*^72^, we utilized the KP3948 strain with the genotype eri-1(mg366) IV; lin-15B(n744) since *eri-1* mutations have been demonstrated to increase the response of neurons to dsRNA^73^. These worms were age synchronized in a similar manner to those fed pET-22b-expressing Rosetta^TM^ 2 cells, with a notable departure following L1 pause where worms were instead plated onto NGM plates containing 100 mM IPTG (dioxane-free, Thermo-Scientific), 100 μg/mL Ampicillin, and 10 μg/mL Tetracycline, and seeded with 250 μL of HT115(DE3) at a concentration of OD_600_ = 5.0. Like the plates used for pET-22b expression, these plates were also incubated at room temperature in the dark at 20°C for 48 h before adding L1 paused worms. After a 30-h incubation period, their behavior was recorded and compared to worms fed the same HT115 (DE3) bacteria producing dsRNA for GFP codon optimized for *O. camponoti-floridani* (pL4440_OcfOptGFP; Data S8).

#### Video recording and computational analysis of C. elegans behavior

For the recording of nematode behavior, an Eo*Sens*® 25CXP High-Speed CMOS Camera was suspended above a plexiglass panel with a 24 VDC inferred LED light (Metaphase Technologies, Inc.) shining upwards towards the camera. A plastic-lined black curtain was draped over the setup to shroud the viewing area in darkness. Approximately ten worms were picked onto each 35 mm petri dish (Greiner) containing 5 mL of NGM media. These plates were then placed onto the plexiglass platform in the viewing area for recording without the lid. After placement, the drape was lowered to block out any light from the experiment and recordings were performed in 6 min intervals following protocols outlined in Beckerson et al.^30^ The recordings were performed in a sequential randomized block format, alternating between randomized control and treatment blocks to account for any temporal, development, temperature, or light exposure effects during picking (Table S3). Analysis of the resulting videos was performed using a custom computational pipeline run with MATLAB (The MathWorks, Inc.)^32^. The individual worm videos cropped from the larger worm arenas are deposited at Zenodo under the DOIs <u>10.5281/zenodo.16760960</u> and the raw data output for each analysis is included in Data S10.

### Behavioral tests in C. floridanus

#### Ant collection

A total of six *C. floridanus* colonies were collected from three separate field sites in Central Florida, USA (Data S11). These collections were permitted by Seminole County Natural Lands. Exact collection location and characterization (i.e., natural nesting substrate and relative colony size) for each of the colonies used for the various behavioral tests can be found in Table S4. Colonies were housed in National Sanitation Foundation certified BPA Free 10 L, 15 x 32.5 x 26.5 cm polypropene Gastronorm containers (Araven) lined with talcum powder (Fisher Scientific) applied dry to the walls to prevent ants from climbing out of the containers. Ants were also provided with makeshift nesting tubes made from ROTILABO^®^ aluminium foil (Carl ROTH)-wrapped 50 mL Falcon tubes (Cellstar) with moisten cotton balls (Etos) at the bottom for humidity. Colonies were fed autoclaved crickets (Size 4, ∼7 mm) twice a week and provided with 10 mL glass tubes of tap water and 10% sucrose dissolved in Milli-Q water plugged with cotton (Etos) *ad libitum*. Prior to the study, these housing boxes were kept at room temperature (∼25°C) on a shelving unit with a 12 h/12 h day-night cycle facilitated by LED strips (LedKoning; 120 leds/meter, IP65, 24 Volt, 5500K) on a timer.

#### dsRNA production

The *C. floridanus* genome includes a *sid-1* orthologue^74^ that presumably facilitates systemic RNAi through the same passive bi-directional dsRNA transport system as *C. elegans*^75^. However, the ideal fragment size for RNA-mediated knockdown in insects differs from nematodes, ranging from 140-520 nucleotides^76^. In *C. floridanus*, a fragment size of around 400 bp has been previously shown to result in successful knock down^74^ and can be introduced via injection^77^. The dsRNA fragment was designed using the NCBI primer blast tool with default settings. The product size was set from 200-400 bp and the organism was set to Camponotus floridanus (taxid: 104421). The primer pair was selected from candidates with a GC content of 40-60%, a GC-clamp, self-complementary values < 7, 3’ complementary values < 4, and without any homopolymer stretches longer than three nucleotides and synthesized by Biolegio (Nijmegen, NL). We also screened for potential 3’ and self-annealing hairpin formation using the Oligonucleotide Properties Calculator (v3.10). This resulted in the selection of a dsRNA fragment 206 bp in length (PS1 dsRNA sequence in Data S8). This fragment matches all three transcript variants of PS1. The fragment was amplified with Q5® High-Fidelity DNA Polymerase (New England Biolabs) from ant cDNA using the primers pL4440_PS1 Fw and pL4440_PS1 Rv (Data S7) and cloned into the same pL4440 vector used to knock down *scrm-1* in *C. elegans*. Cloning was performed via Gibson Overlap PCR using the NEBuilder® HiFi DNA Assembly Cloning Kit (New England Biolabs) and pL4440 linearized with EcoRV (New England Biolabs). The resulting plasmid was named pL4440_PS1 (Data S8).

To isolate the dsRNA for injection into ants, HT115 strains were transformed with pL4400_PS1 to produce dsRNA for PS1 and pL4440_OcfOptGFP to produce dsRNA for GFP to use as a control. Transformation followed conventional heat shock transformation protocols with CaCl_2_-treated bacteria and was validated through colony PCR with the primers pL4440 Screen Fw & Rv (Data S7). Resulting transformants were shaken overnight at 200 rpm in 5 mL of LB + 100 μg/mL Amp + 10 μg/mL Tet liquid media at 37°C. The next morning, the OD_600_ of each tube was checked and 500 μL was transferred to 50 mL of TB + 100 μg/mL Amp + 10 μg/mL Tet in a 500 mL flask. The flask was shaken at 200 rpm in a 37°C incubator until it reached an OD_600_ = 0.4, after which IPTG (dioxane-free, Thermo-Scientific) was added to the flask to a final concentration of 100 mM and the flask was moved to a 20°C shaker at 180 rpm and shaken overnight. The culture was then transferred to 50 mL Flacon tubes for centrifugation at 5,000 x g for 5 min to pellet the cells. Supernatant was removed and the cells were resuspended in 800 μL of PBS + 0.1% SDS and transferred to 2 mL microcentrifuge tubes for extraction. dsRNA was extracted by vortexing each tube with 800 μL of Trizol and incubating at room temperature for 5 min. The extracted dsRNA was then isolated using alcohol precipitation and treated with Turbo^TM^ DNaseI and RNase A before purifying with PCI 25:24:1. Resulting dsRNA extracts were resuspended in PBS to a concentration of ∼2.5 μg/μL for injection.

#### SBP production

Periplasmic extracts of *O. camponoti-floridani* SBP were produced using the same Rosetta^TM^ 2 strain used for the feeding exposure tests in *C. elegans*. Because the pET-22b plasmids share the same DE3 background as the pL4440 plasmids, protein production was performed using nearly the same protocol as the aforementioned dsRNA preparation. Cells harboring the pL4440_SBP and pL4440_OcfOptGFP plasmids were streaked onto LB agar plates supplemented with 100 μg/mL ampicillin and 25 μg/mL chloramphenicol (Sigma-Aldrich, Germany) and incubated overnight at 37 °C. The following day, single colonies were used to inoculate 5 mL LB medium with the same antibiotics and grown overnight at 37 °C, shaking at 200 rpm. The resulting suspensions were then used to inoculate a 50 mL starter culture at an OD_600_ of 0.2 in Terrific Broth (0.6 g/L tryptone, 1.2 g/L yeast extract, 43.45 mz glycerol, 27 mM potassium phosphate dibasic, and 8.67 mM potassium phosphate monobasic) supplemented with 100 μg/mL ampicillin and 25 μg/mL chloramphenicol. These cultures were incubated at 37 °C with shaking at 200 rpm until they reached an OD_600_ of 0.8–1.0, at which point protein expression was induced by adding IPTG (dioxane-free, Thermo-Scientific) to a concentration of 1 mM. The temperature was lowered to 28°C to favor protein production and incubated for an additional 2 h, shaking at 200 rpm.

Protein was harvested from the periplasmic space by first pelleting the cells via centrifugation at 3,000 x g for 10 minutes at 4°C. The pellet was carefully resuspended in cold TES buffer (30 mM Tris-HCL, 500 mM sucrose, and 1 mM EDTA at pH 8.0) at 3 mL per 1 g wet cell weight using a wide-bore pipette tip. The new cell suspension was then incubated on ice for 30 minutes with gentle stirring. The outer membrane was then ruptured using 1:1 cold osmotic shock buffer (5 mM MgSO4) and incubated on ice for an additional 10 minutes with gentle stirring. The culture was then centrifuged at 7.500 x g for 10 minutes at 4°C to pellet the spheroplasts and collect the supernatant containing the periplasmic protein. The protein was concentrated using ultracentrifugation by loading the periplasmic fraction onto 15 mL 3.000 MWCO Amicon® Ultra Centrifugal Filters and centrifugating for 30 minutes at 4.000 x g in a swinging rotor at 4°C. The supernatant containing the protein was resuspended and used for protein visualization and injections.

#### Microinjection of C. floridanus with dsRNA and SBP

Injections were performed using borosilicate glass micropipettes (Fisher) pulled with a glass puller (model: PC-100, Narishige) configured to use a single heating step at 62°C with all weights. To prepare the ants for injection with dsRNA, we first synchronized their circadian rhythm by transferring groups of approximately 20 ants from their colony housing to new housing containers (Denox) lined with talcum powder (Fisher Scientific). These containers were also fitted with a 2 mL microcentrifuge tube (Greiner) of 10% sucrose, a 2 mL microcentrifuge tube (Greiner) of tap water plugged with cotton balls (Etos), and a small housing unit created by cutting a small hole in an 89 mm Fisherbrand^TM^ Hexagonal Antistatic Weighing Boat (Fisher) lined with ROTILABO^®^ aluminium foil (Carl ROTH). Once acclimated, these housing units were moved to a Micro Clima-Series™ Economic Lux Chamber (Snijders Labs) under constant light with a temperature of 25°C and 70% humidity for 2 days. After 2 days, the parameters were changed to a 12 hr day/night cycle with a temperature of 25°C with 70% humidity during the day and a temperature of 20°C with 70% humidity during the night. 1 day prior to injection, ants were starved for 24 h by removing the tubes containing the sugar and water. After 24 h, these containers were moved to the lab bench in the light at room temperature for injection. SBP injections were performed in a similar manner but did not separate ants from colony until injection, provided food and water in 15 mL tubes, and were provided larger housing units prepared from 50 mL Falcon tubes wrapped in aluminum foil.

All ants were injected with 0.5 μL using the technique outlined in Beckerson et al.^46^ Following injection, ants were placed into open-air viewing arenas and briefly observed for any signs of physical trauma from the injection process that would interfere with normal ant movement (e.g., paralysis or disablement of any limbs). Any ants that exhibited signs of trauma were immediately anesthetized using a petri dish (Cellstar) on ice and later euthanized by moving the petri dish to –20°C for 48 h. Importantly, any arenas that held a traumatized ant were also promptly discarded and a new arena was used for the next injection to prevent any effects from potential pheromones left behind by the prior occupants. If the ants did not show signs of trauma, we moved them to a viewing arena corresponding to their respective experiments.

#### Video recording of C. floridanus

All the videos for each set of tests were recorded using a randomized block format to mitigate any temporal variables that might affect behavior (Table S5). For the antennation tests, arenas were prepared by dry brushing talcum powder (Fisher Scientific) onto the walls of “tall-wall” 60 mm x 20 mm petri dish (Cellstar), which preliminary studies showed was sufficient to prevent escape of the ant without a lid, allowing us to improve video quality by reducing reflection and glare. A piece of uncoated, woodfree computer paper cut to the dimensions of the arena was then fastened to the inside bottom of the dish with two strips of double-sided adhesive roller tape (Pritt) to provide a graspable surface and help prevent ant slipping due to the talcum powder. The paper also further reduced reflection and glare, providing better contrast between the ant and the arena background. Just prior to injection, ants were randomly painted with either silver or black non-toxic water-based paint (POSCA; Table S5). This was done to help keep track of which ant was injected with either the treatment or control in a random way that is not detectable during manual scoring of the resulting videos. Ants were then paired two per arena and experiments were recorded in the light at 60 fps for 10 min using two or three different infrared GoPro Hero6s (GoPro, Inc.) in a rolling fashion.

For the aversion assays and preceding nest behavior analyses, injected ants were first transferred to a new National Sanitation Foundation certified BPA Free 1.1 L, 6.5 x 17.6 x 16.2 cm polypropene Gastronox containers (Denox) lined with talcum powder (Fisher Scientific) lined with the same computer paper used in the antennation tests. For ants injected with dsRNA, each arena was provided with the same shelter and 2 mL microcentrifuge tubes (Greiner) of 10% sucrose and tap water as their previous container and returned to the Micro Clima-Series™ Economic Lux Chamber (Snijders Labs) for 36 h. After their 36-h incubation, the ants were then moved to the viewing area in complete darkness for group behavior observations. These recordings started ZT 9-12, which marks the beginning of their natural foraging period. A 2 x 2 cm piece of Whatman paper cut from a 46 x 57 cm sheet (VWR) spotted with 50 μL of citronella oil (De Tuinen) and placed into the corner of the arena and recordings were started. The videos were captured in high definition at 30 FPS for 10 min using an infrared USB 2.0 Camera HD (ELP) and the Movavi Screen Recorder (v23.1.0).

For ants injected with SBP extracts, no shelter, food, or water was provided. Instead, the arenas were moved directly to the viewing area in the light for group behavior observations. We performed these tests during the day because the injections required light to perform. During the recordings, ants were allowed to acclimate for 10 min to allow the effects of SBP to take hold before the citronella pads were added to the arena. This duration was determined by performing a small pilot test to screen for behavioral effects of SBP injection after a 10, 20, and 30-min acclimation period. After adding the pad, the recordings continued for an additional 10 min. Videos were recorded using the same USB 2.0 Camera HD (ELP) as the RNAi tests. Resulting videos were processed into 10 min clips immediately following addition of the citronella pads for the aversion test analyses and 5 min clips directly prior to addition of the citronella pads for nest behavior analyses. The later clips were shorter to ensure that even the most recently injected ants had at least 5 min of SBP exposure before video analysis. Post-processing was done using Movavi Video Editor Plus (v22.4.1). All of the processed videos used in the analysis phase of this study have been deposited at Zenodo under the DOI <u>10.5281/zenodo.16760416</u>.

#### Manual scoring and computational analysis of C. floridanus behavior

The antennation tests were scored manually by counting the number of antennation events, examples of which are provided in Fig. 5B. This scoring was done in a blinded fashion with each video being assigned a randomized 2 Greek letter identifier viewing. After scoring, the metadata for each experiment was used to unblind the results for further analysis. The aversion tests and the nest behavior tests were analyzed using FIJI and trackmate. We converted each 30-fps video to AVI format, keeping every 15th frame (0.5 sec) to help avoid oversampling while still capturing dynamic behaviors. To assess the spatio-temporal distribution of ants in response to the addition of a citronella pad, we used a combination of Trackmate^78^ in FIJI and Python^79^. Once imported into FIJI, we segmented ants using the threshold option in trackmate^78^ (threshold = 182) and applied filters based on spot size and quality to exclude erroneous spots. We then calculated the distance of every ant on every frame from the center of the citronella pad on that frame. A similar approach was also used for the nest behavior tests. After framerate decimation and ant segmentation through FIJI, we calculated the nearest-neighbor distance for every ant on every frame and reported an average of these values for each frame. We favored this approach over clustering algorithms due to the scarcity of data points on each frame to inform clustering cutoffs and because nearest neighbor measurements give a metric that combines both group clustering and individual interactions. Over the course of the videos, ants that cluster or spend time more closely interacting show a lower overall average of nearest-neighbor distance compared to ants which have shorter interactions or more diffuse and unstable groups. The raw data output for each of these tests is included in Data S12, subcategorized by experiment in subfolders.

### Statistical analyses

All statistical analyses were performed using R and R-studio (Version 2023.06.0+421). The statistical tests, packages used, and R code are provided in Data S6.

## Supplementary Text

**Sup. Text 1. Colorblind-friendly code for PyMOL**

show cartoon

bg_color black

set antialias=1

set orthoscopic=1

set gamma=1.15

set cartoon_fancy_helices, 1

set cartoon_fancy_sheets, 1

set_color SunnyOrange, [255,194,10]

set_color BrightSkyBlue, [0,202,255]

set_color AbsoluteRed, [255,0,0]

set_color Megaviolet, [189,3,186]

set_color PlainBlack, [0,0,0]

set_color PlainYellow, [255,255,0]

set_color PlainWhite, [255,255,255]

set ray_shadows, 0

set ray_trance_fog, 1 util.cbac

spectrum b, white **###name of corresponding set_color here###**

## Supplementary Figures

**fig. S1.**
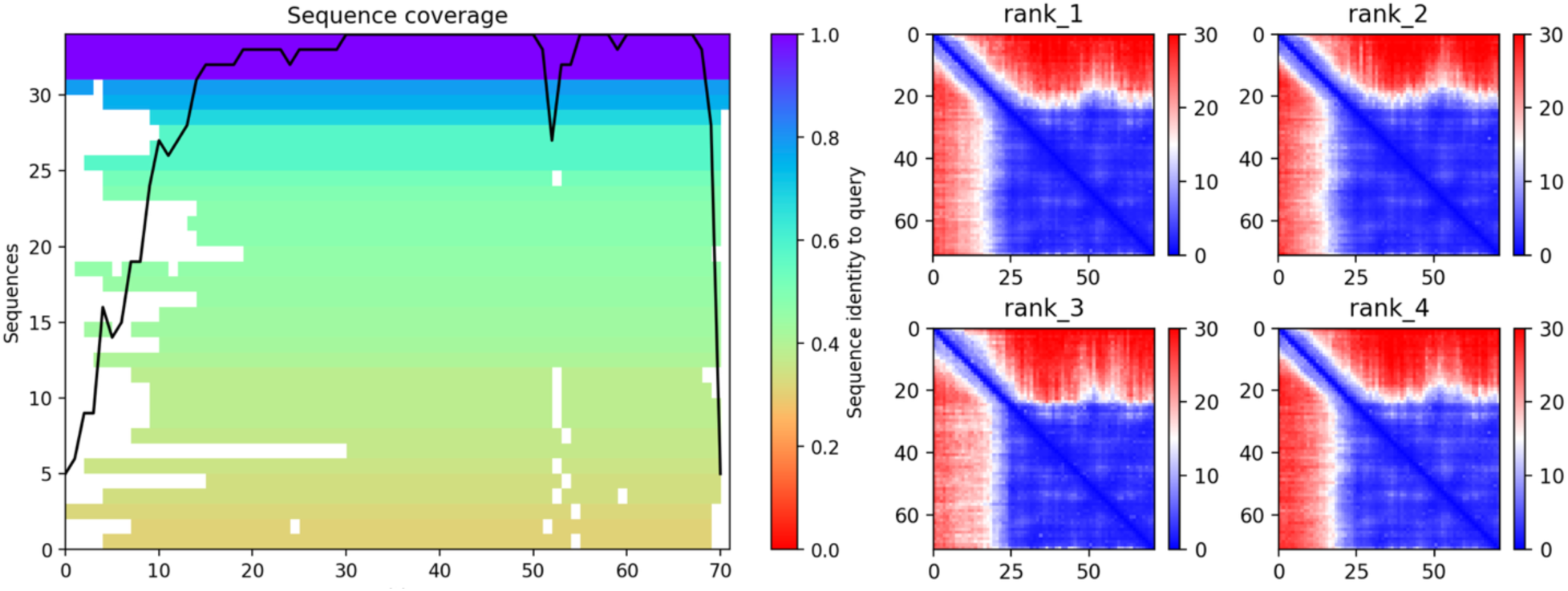
Sequence coverage and Predicted Aligned Error (PAE) scores for Ophcf2|06345. Multiple sequence alignments for the Ophcf2|06345 peptide from AlphaFold2 v2.3.1 with their level of sequence identity, left. The top four highest-ranked models for PAE plots, right. Both results reflect the scores for the entire length of the Ophcf2|06345 peptide sequence, including both the signal peptide region and the mature protein region. The low confidence regions illustrated in red are found within the signal peptide region. The mature peptide region shows a high degree of certainty for the predicted model (Fig. 1; Data S3).

**fig. S2.**
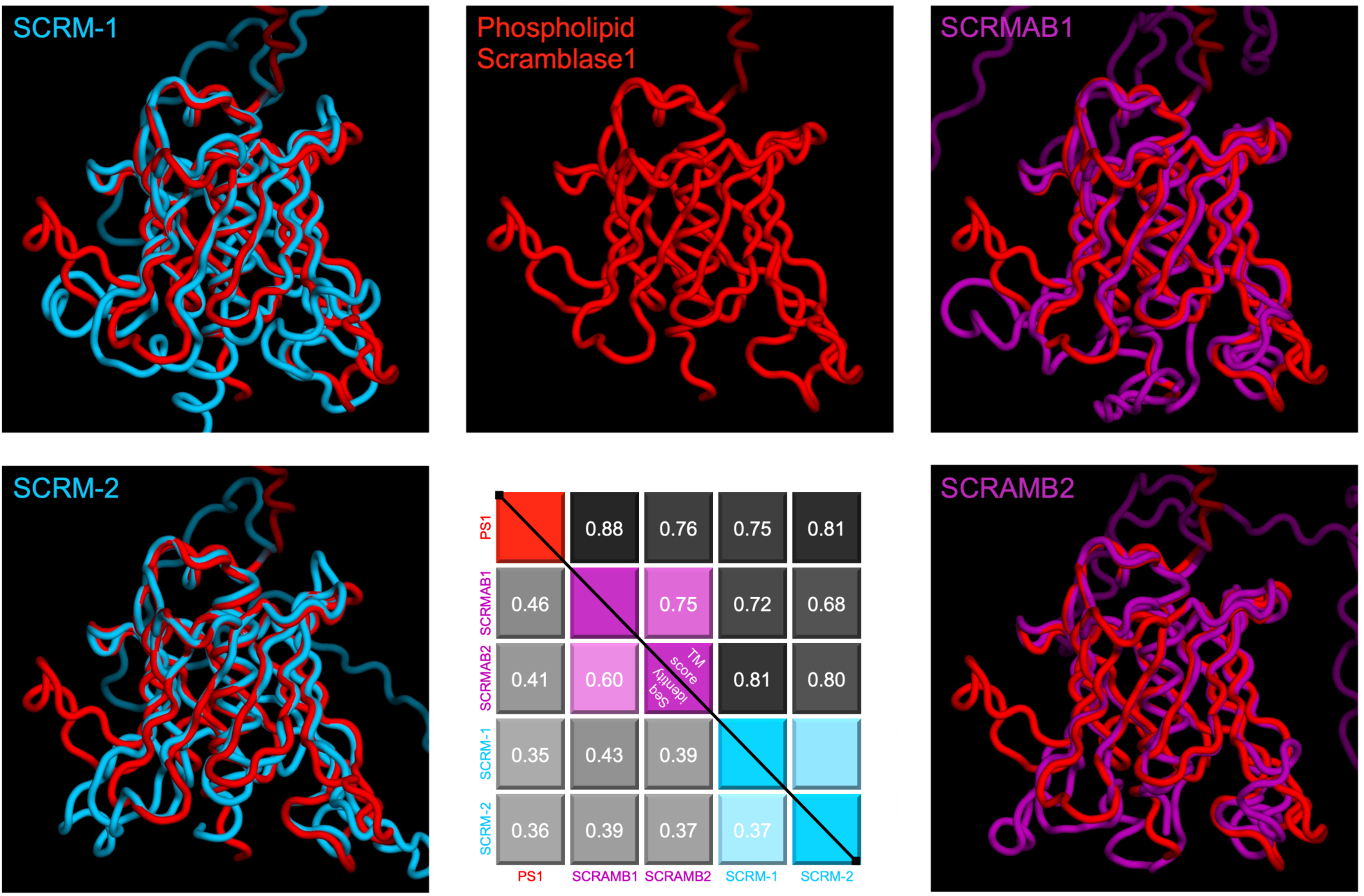
Computational modeling and structural comparison of Scramblase homologs. The AlphaFold2-predicted models for *C. floridanus* Phospholipid Scramblase 1 (PS1), red, its Scramblase proteins 1 and 2 homologs in *C. elegans* (SCRM-1 and SCRM-2), blue, and *D. melanogaster* (SCRAMB1 and SCRAMB2), purple. Superimpositions of PS1 with these homologs demonstrate a high degree of structural similarity between the homologs. TM scores for each of these comparisons are shown in the top right of the matrix chart with corresponding amino acid sequence identities for each comparison shown in the bottom left.

**fig. S3.**
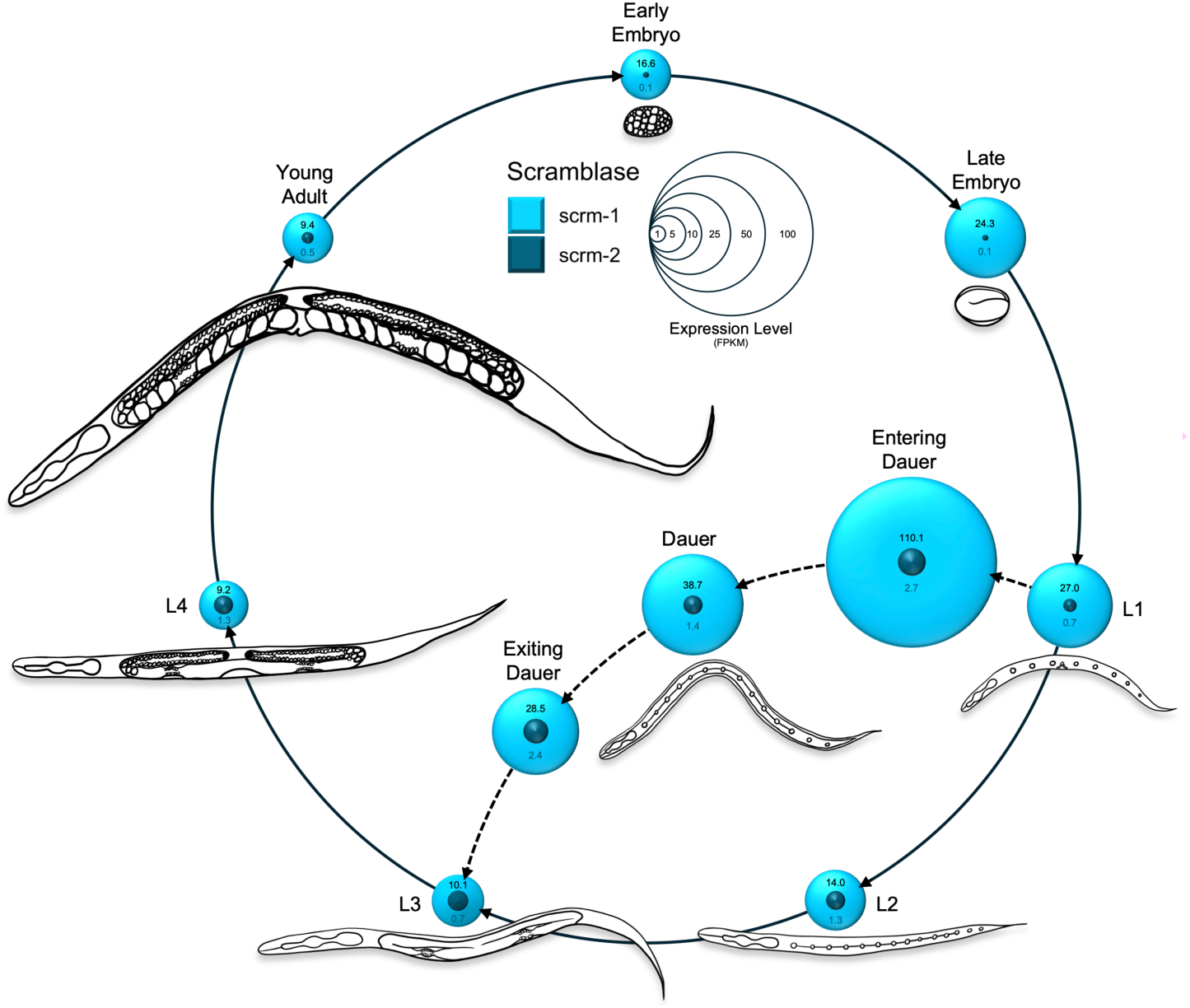
Expression levels for SCRM-1 and SCRM-2 across the *C. elegans* life cycle. Expression levels for Scramblase-1 and Scramblase-2 are represented by spherical modules. The diameter of these spheres is based on the FPKM values for each stage of nematode development collected from WormBase^80^. Images of each life cycle stage were drawn using the Procreate app.

**fig. S4.**
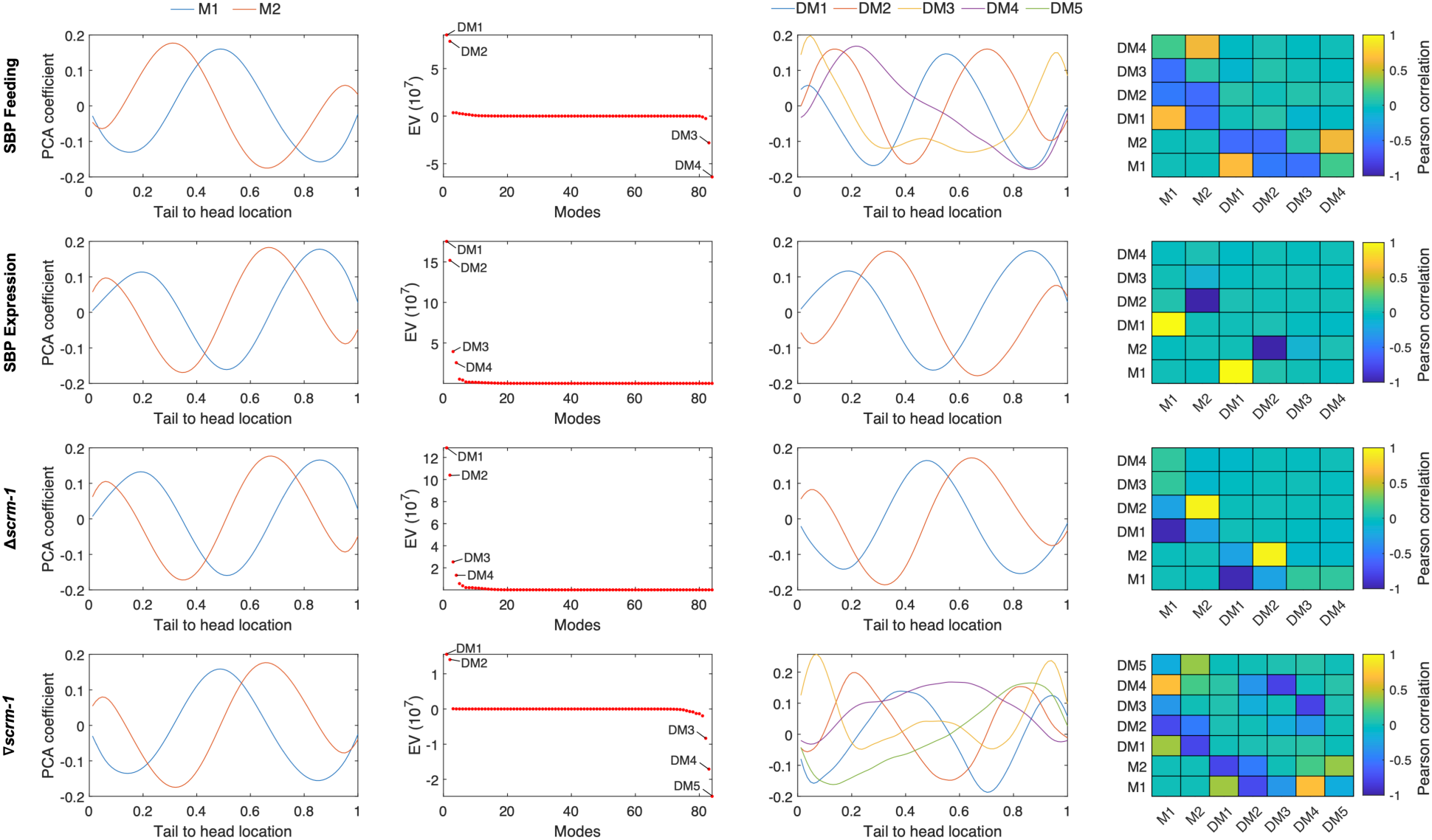
Posture analysis during C. elegans crawling using PCA and differential eigenvalue spectrum analysis. The data is organized into rows based on the experimental parameters of Fig. 4. (A) A Principal components analysis (PCA) is performed separately for each experiment. The curvature along the worm body is decomposed into PCA coefficients, sometimes referred to as “eigenworms”, with a respective amplitude (Fig. 4). The first two PCA coefficients with the largest eigenvalues look sinusoidal. When multiplied by the respective time-varying amplitudes, they yield the undulatory body wave observed in *C. elegans* locomotion. (B) Eigenvalue spectrum of the difference covariance matrix Cov_pert – Cov_ctrl. Outlier differential modes (DM), for which the variance is much larger in either the perturbation or the control, are labelled and further analyzed. (C) The coefficients of the identified DMs are similar to the first two PCA modes. (D) The absolute Pearson correlation values between the most important DMs and the first two PCA modes indicate that the difference in shape between perturbation and control arises from a difference in a curvature amplitude (i.e. a worm bends more strongly) rather than a difference in shape.

**fig. S5.**
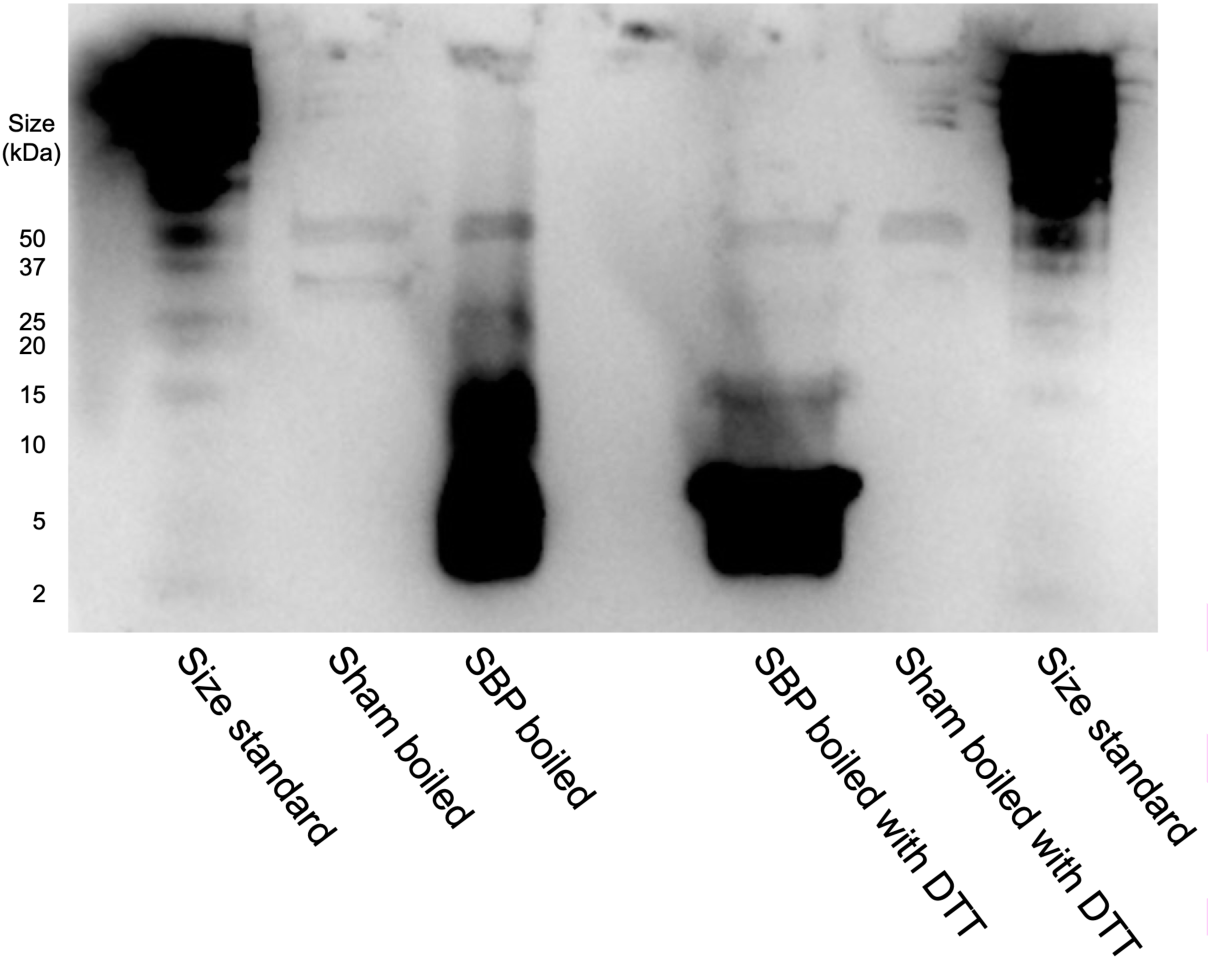
Evidence of disulfide-bond formation in the scramblase binding peptide (SBP). Western blot of periplasmic protein samples expressing SBP fused to a C-terminal 6XHis-tag under reducing dithiothreitol (DTT) and non-reducing conditions. Under non-reducing conditions, SBP exhibits slower migration on SDS-PAGE, whereas reduction with DTT demonstrates faster migration, consistent with the formation of intramolecular disulfide bonds. SBP is expected at 7.64 kD size. Boiled bacterial lysates are shown in lanes 2 and 3. Boiled lysates with the addition of DTT are shown in lanes 5 and 6. Lanes 1 and 7 contain size standards with units displayed in line with their corresponding bands to the left of the figure.

**fig. S6.**
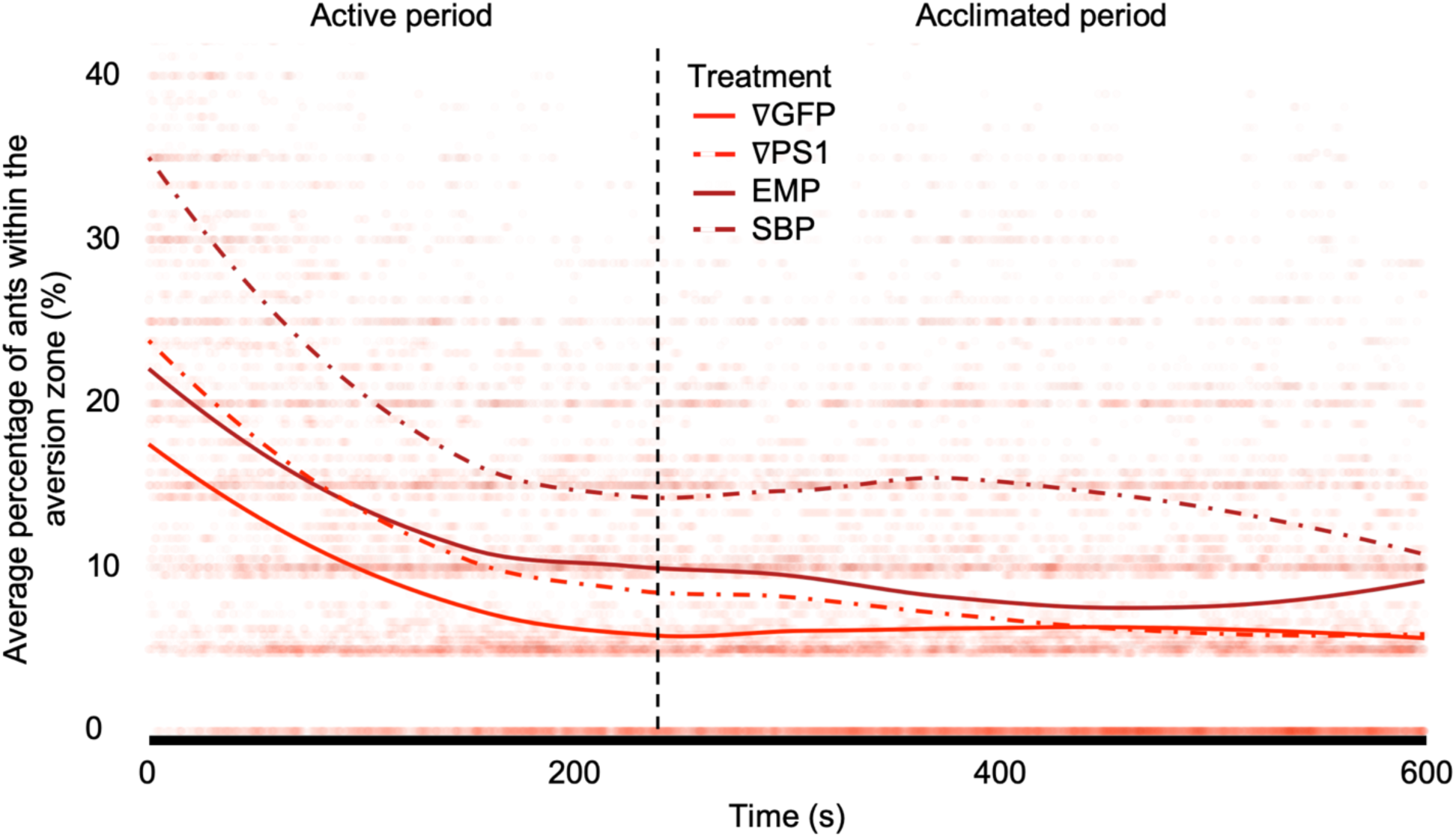
Aversion zone proximity of ants during exposure to citronella oil. The percentage of ants within the aversion zone (4 cm from the center of the citronella pad) every 0.5 sec across a 10 min recording. Each data point is shown with a high degree of transparency in the background. The first 4 min of recording depict an active response to the addition of a foreign object into the nest arena. After 4 min, the ants settle down with a majority of them positioning themselves away from the citronella pad. A small portion of the group, ∼25%, continues with exploratory behaviors resulting in re-exploration of the pad in the final 6 min of the video.

**fig. S7.**
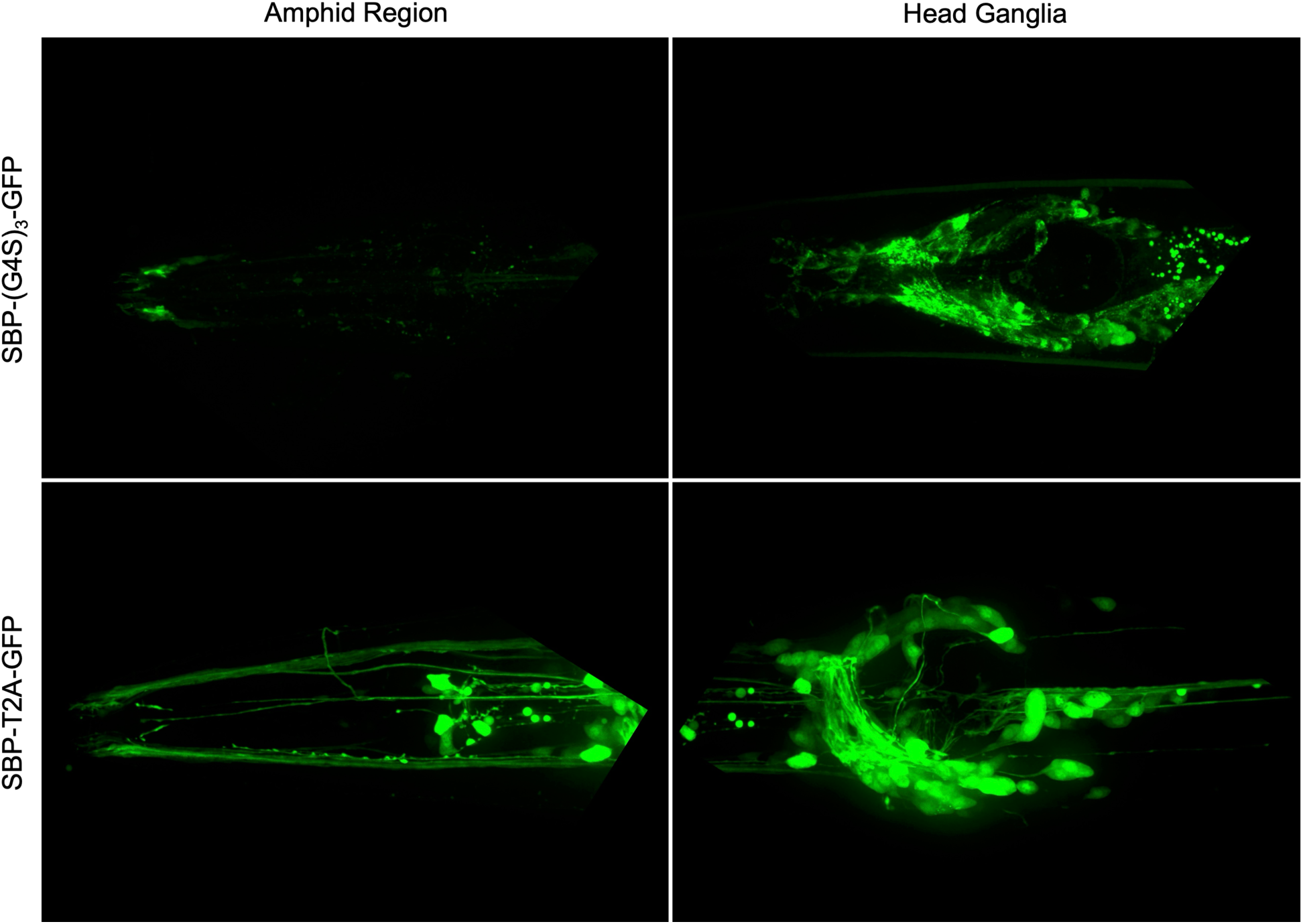
Expression profiles of GFP in *C. elegans*. Spinning disc confocal imagery of the *C. elegans* amphid region, left, and nerve ring, right, captured with a 491 nm laser to excite the GFP fluorophore, shown in green. The images in the top row depict worms expressing *pPrab-3_CelOptSBP-(G4S)3-GFP_Tlet-858*, which codes for secreted GFP tethered to SBP with a (G4S)_3_ linker. The images on the bottom row depict worms expressing pPrab-3_CelOptSBP-T2A-GFP_Tlet-858, which codes for polycistronic expression of GFP fluorophore separated from the secreted SBP via ribosomal skipping facilitated by a T2A linker. The brightness is standardized between the left and right image for each row, but not between each row. This is due to the sheer difference in brightness of the bottom row images, which can be noted by the lack of visible autofluorescence anterior to the nerve right in the bottom right picture. The brightness of the worms utilizing the T2A linker was high enough for selection by eye under excitation conditions, while the brightness of the (G4S)_3_ line was too low for selection, requiring the use of a helper plasmid expressing *rol-6*(su1006) during transformation.

**fig. S8.**
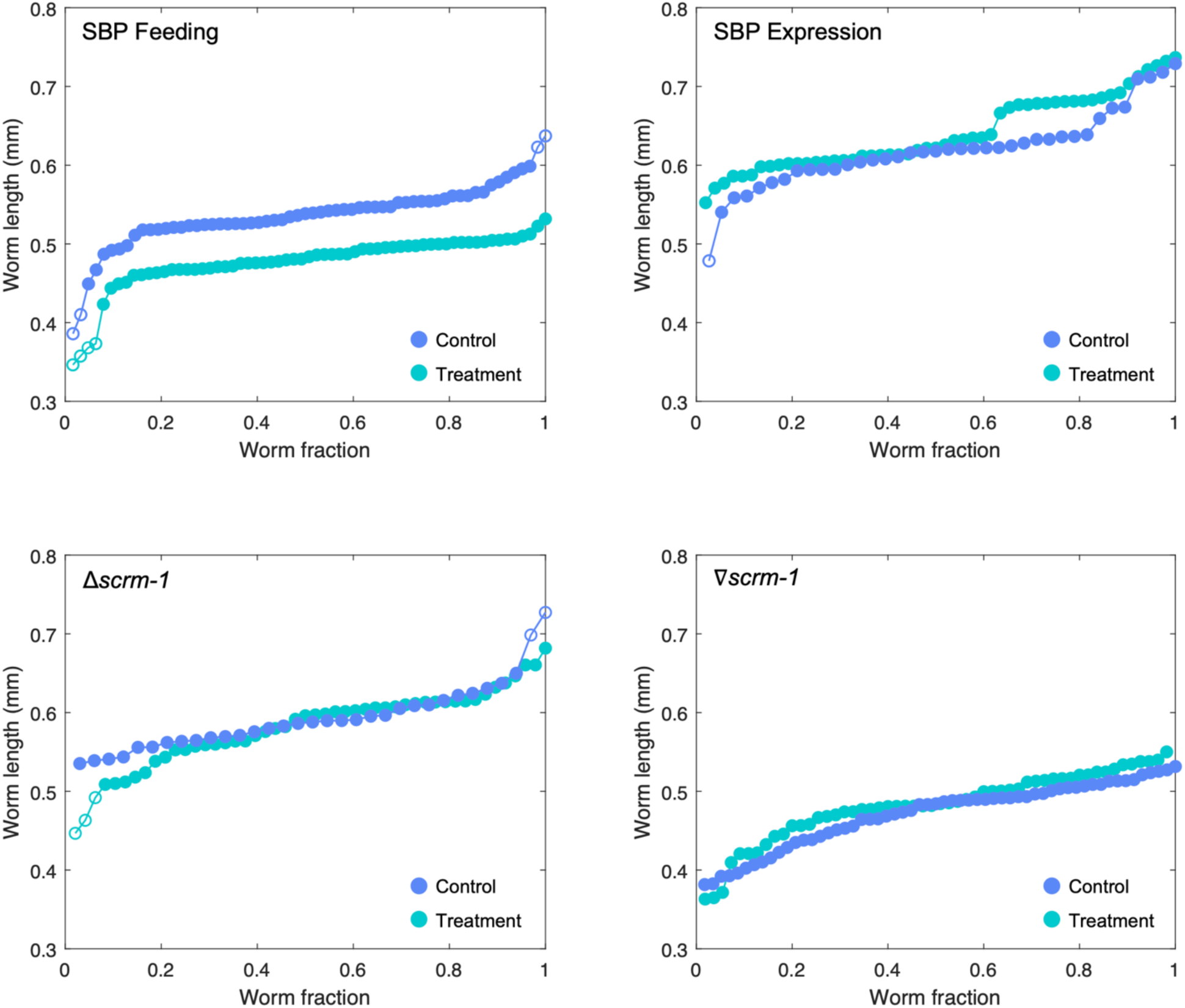
Size standardization of worms per behavioral experiment. Graphs depicting the length of each worm used in this study, separated by experiment and arranged left to right from smallest to largest. While the duration of each age standardization process used prior to video recording in each experiment was the same, strains and growth conditions used between each experiment were different. Importantly, these variables were standardized between test and control groups within each experiment, allowing for robust comparison to be made for each experimental variable, but not between experiments. The different growth variables resulted in worms of slightly different length depending on the experiment; however, very similar size-distributions for the control and treatment groups were obtainable within each experiment. Any outliers were removed from their respective analyses (14 in total) and are indicated in the graphs using empty fill circles.

## Supplementary Tables

**Table S1.** Colorblind friendly palette for scientific figures.

| Color Name | True | Prot. | Deut. | Trit. |
| --- | --- | --- | --- | --- |
| Sunny Orange | #FFC20A |  |  |  |
| Bright Sky Blue | #00CAFF |  |  |  |
| Absolute Red | #FF0000 |  |  |  |
| True Black | #000000 |  |  |  |
| Bright Yellow | #FFFF00 |  |  |  |
| Mega Magenta | #BD03BA |  |  |  |

**Table S2.**
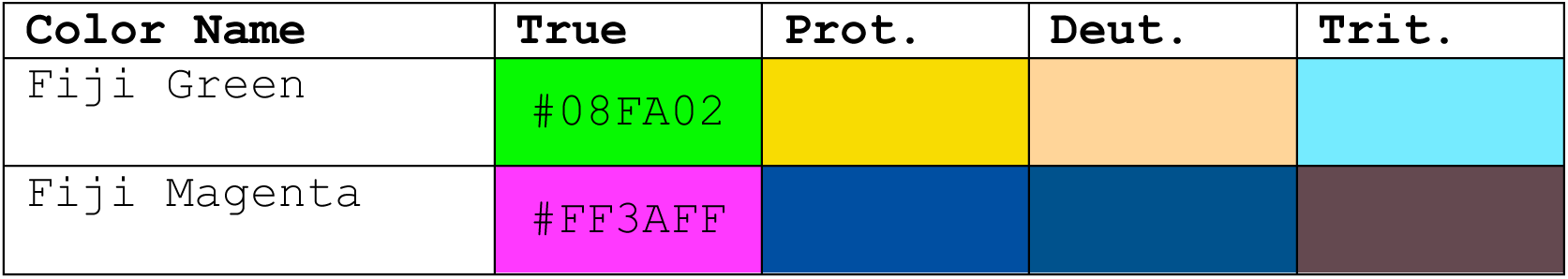
Colorblind friendly palette for colocalization imaging.

| Color Name | True | Prot. | Deut. | Trit. |
| --- | --- | --- | --- | --- |
| Fiji Green | #08FA02 |  |  |  |
| Fiji Magenta | #FF3AFF |  |  |  |

**Table S3.**
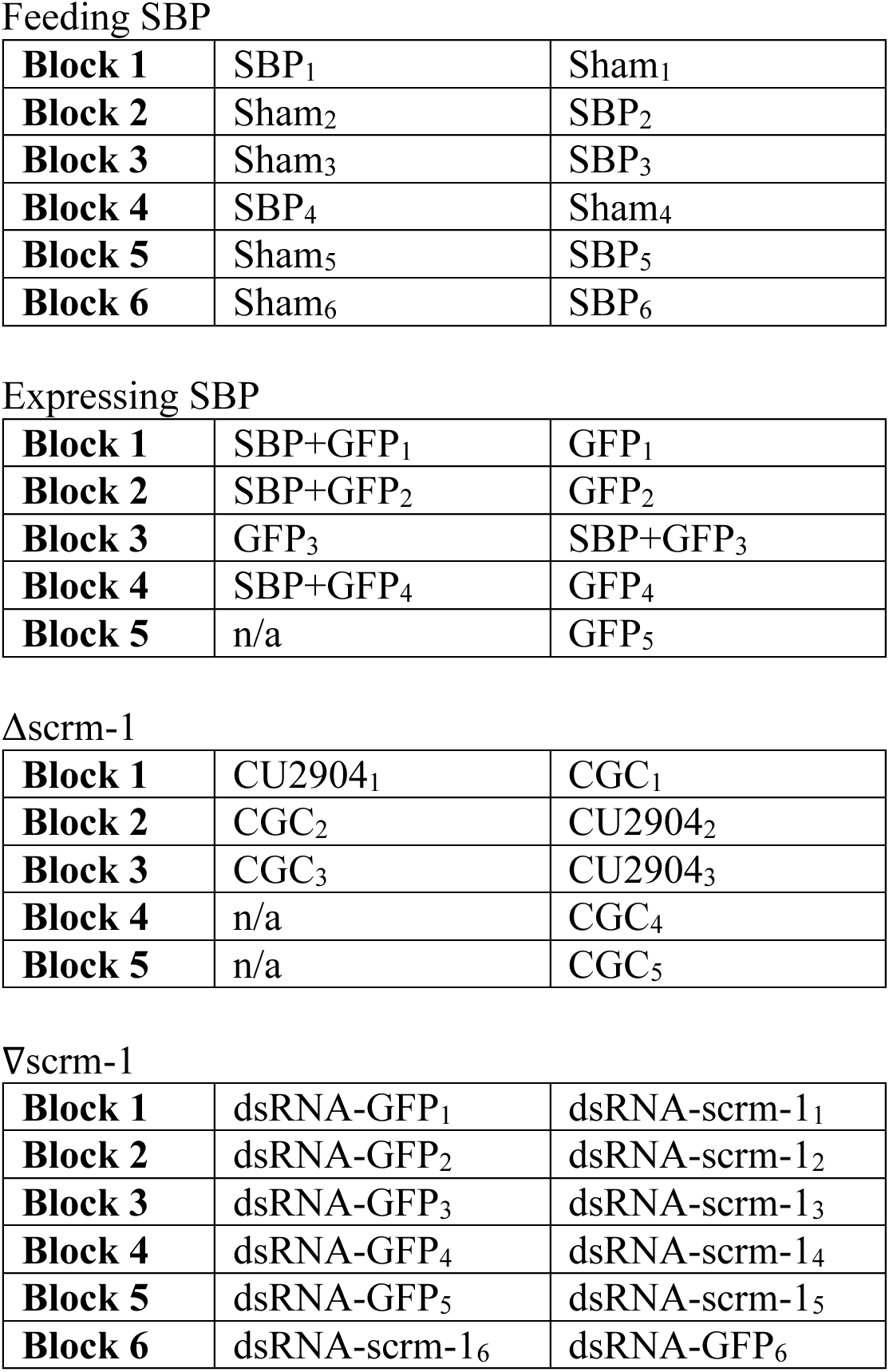
The randomized block format for recording worm behavior.

**Table S4.** Ant colony information.

| Name | Wilderness Area | Substrate | Size* | Coordinates |
| --- | --- | --- | --- | --- |
| 2024Chul5 | Chuluota | Log | 16 tubes | 28°37'10.82" N, 81°3'44.02" W |
| 2024Gen17 | Geneva | Log/Ground | 5 tubes | 28°42'18.83" N, 81°7'31.23" W |
| 2024LH15 | Lake Harney | Tree stump | 7 tubes | 28°47'16.56" N, 81°3'52.35" W |
| 2024LH11 | Lake Harney | Log | 8 tubes | 28° 47' 15.03" N, 81° 3' 55.03" W |
| 2024Gen19 | Geneva | Tree stump | 5 tubes | 28° 42' 6.18" N, 81° 7' 30.63" W |
| 2024LH14 | Lake Harney | Tree stump | 8 tubes | 28°47'15.03" N, 81°3'55.03" W |

\*Ant colony size was determined by allowing ants to self-assort into 50 mL Falcon tubes.

**Table S5.**
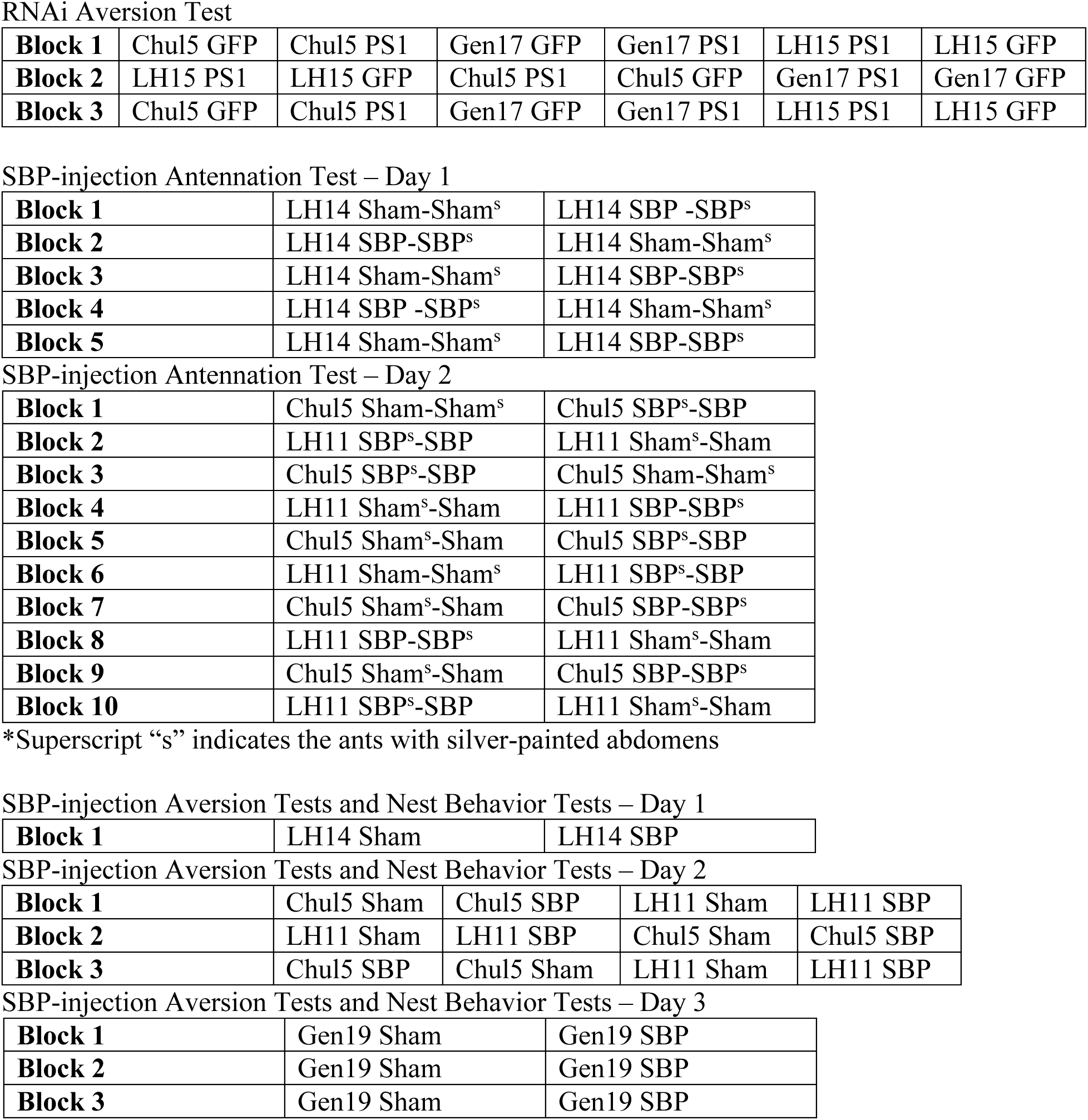
The randomized block format for recording ant behavior.

RNAi Aversion Test
|  |  |  |  |  |  |  |
| --- | --- | --- | --- | --- | --- | --- |
| **Block 1** | Chul5 GFP | Chul5 PS1 | Gen17 GFP | Gen17 PS1 | LH15 PS1 | LH15 GFP |
| **Block 2** | LH15 PS1 | LH15 GFP | Chul5 PS1 | Chul5 GFP | Gen17 PS1 | Gen17 GFP |
| **Block 3** | Chul5 GFP | Chul5 PS1 | Gen17 GFP | Gen17 PS1 | LH15 PS1 | LH15 GFP |

SBP-injection Antennation Test – Day 1
|  |  |  |
| --- | --- | --- |
| **Block 1** | LH14 Sham-Sham<sup>s</sup> | LH14 SBP-SBP<sup>s</sup> |
| **Block 2** | LH14 SBP-SBP<sup>s</sup> | LH14 Sham-Sham<sup>s</sup> |
| **Block 3** | LH14 Sham-Sham<sup>s</sup> | LH14 SBP-SBP<sup>s</sup> |
| **Block 4** | LH14 SBP-SBP<sup>s</sup> | LH14 Sham-Sham<sup>s</sup> |
| **Block 5** | LH14 Sham-Sham<sup>s</sup> | LH14 SBP-SBP<sup>s</sup> |

SBP-injection Antennation Test – Day 2
|  |  |  |
| --- | --- | --- |
| **Block 1** | Chul5 Sham-Sham<sup>s</sup> | Chul5 SBP<sup>s</sup>-SBP |
| **Block 2** | LH11 SBP<sup>s</sup>-SBP | LH11 Sham<sup>s</sup>-Sham |
| **Block 3** | Chul5 SBP<sup>s</sup>-SBP | Chul5 Sham-Sham<sup>s</sup> |
| **Block 4** | LH11 Sham<sup>s</sup>-Sham | LH11 SBP-SBP<sup>s</sup> |
| **Block 5** | Chul5 Sham<sup>s</sup>-Sham | Chul5 SBP<sup>s</sup>-SBP |
| **Block 6** | LH11 Sham-Sham<sup>s</sup> | LH11 SBP<sup>s</sup>-SBP |
| **Block 7** | Chul5 Sham<sup>s</sup>-Sham | Chul5 SBP-SBP<sup>s</sup> |
| **Block 8** | LH11 SBP-SBP<sup>s</sup> | LH11 Sham<sup>s</sup>-Sham |
| **Block 9** | Chul5 Sham<sup>s</sup>-Sham | Chul5 SBP-SBP<sup>s</sup> |
| **Block 10** | LH11 SBP<sup>s</sup>-SBP | LH11 Sham<sup>s</sup>-Sham |

\*Superscript “s” indicates the ants with silver-painted abdomens

SBP-injection Aversion Tests and Nest Behavior Tests – Day 1
|  |  |  |
| --- | --- | --- |
| **Block 1** | LH14 Sham | LH14 SBP |

SBP-injection Aversion Tests and Nest Behavior Tests – Day 2
|  |  |  |  |  |
| --- | --- | --- | --- | --- |
| **Block 1** | Chul5 Sham | Chul5 SBP | LH11 Sham | LH11 SBP |
| **Block 2** | LH11 Sham | LH11 SBP | Chul5 Sham | Chul5 SBP |
| **Block 3** | Chul5 SBP | Chul5 Sham | LH11 Sham | LH11 SBP |

SBP-injection Aversion Tests and Nest Behavior Tests – Day 3
|  |  |  |
| --- | --- | --- |
| **Block 1** | Gen19 Sham | Gen19 SBP |
| **Block 2** | Gen19 Sham | Gen19 SBP |
| **Block 3** | Gen19 Sham | Gen19 SBP |

## Supplementary Movies

**Movie S1. Z-stack imaging of SBP/SCRM-1 colocalization in the amphid region.** A series of images captured at different focal planes along the z-axis of transgenic *scrm-* 1(he439[*mKate2::scrm-*1]) nematodes expressing pPrab-3_CelOptSBP-(G4S)3-GFP_Tlet-858. GFP-tagged SBP is shown in green and mKate2-tagged SCRM-1 is shown in magenta.

**Movie S2. Z-stack imaging of SBP/SCRM-1 colocalization in the head ganglia.** A series of images captured at different focal planes along the z-axis of transgenic *scrm-* 1(he439[*mKate2::scrm-*1]) nematodes expressing pPrab-3_CelOptSBP-(G4S)3-GFP_Tlet-858. GFP-tagged SBP is shown in green and mKate2-tagged SCRM-1 is shown in magenta.

**Movie S3. Median worm behaviors.** A composite video of worm behavior for the median worm, based on PCA diameter, from each test group as seen by the our computational analysis pipeline. The worm IDs for each panel are SBP Feeding Control Worm 11 and Treatment Worm 4, SBP Expression Control Worm 3 and Treatment Worm 4, SCRM-1 Knockout Control Worm 10 and Treatment Worm 1, and SCRM-1 Knockdown Control Worm 3 and Treatment Worm 4. A midline for each worm is generated in red and superimposed onto each worm. All instances of turning and worm collisions have been removed and appear as black screens for the duration of these events in the video.

**Movie S4. Nestmate Fighting Behavior.** Video example for nestmates being attacked upon placement into arenas by ants exposed to SBP approximately 5-10 min prior.

## Supplementary Data

**Data S1. Bioinformatic pipeline.xlsx**

**Data S2. Secretion signal predictions for SBP homologs.xlsx**

**Data S3. SBP cystine lattice structure.pse**

**Data S4. Y2H assay results.xlsx**

**Data S5. PS1-SBP polar contacts.pse**

**Data S6. R code.R**

**Data S7. Primer list.xlsx**

**Data S8. Plasmid maps.zip**

**Data S9. Codon optimized sequence for SBP expression in nematodes**

**Data S10. Worm behavior tests data.zip**

**Data S11. KML map of ant colonies used in this study.kml Data S12. Ant behavior tests data.zip**

