## Supplementary material for "The First of Us: *Ophiocordyceps* use a novel scramblase-binding peptide to manipulate zombie ants": Data S6

```
##### Worm Code #####
### PCA Plot ###
# Load necessary libraries
library(ggplot2)
library(dplyr)

# Step 1: Filter data by Test and Treatment
plot_data <- Radius_Data %>%
  filter((Test == "Test 1" & Treatment %in% c("pET_emp", "pET_sbp")) |
           (Test == "Test 2" & Treatment %in% c("GFP", "SBP")) |
           (Test == "Test 3" & Treatment %in% c("CGC1", "SCRM1KO")) |
           (Test == "Test 4" & Treatment %in% c("pL4440_gfp",
"pL4440_sbp")))

# Step 2: Set factor levels for consistent ordering
plot_data$Test <- factor(plot_data$Test, levels = c("Test 1", "Test
2", "Test 3", "Test 4"))
plot_data$Treatment <- factor(plot_data$Treatment, levels = c(
  "pET_emp", "pET_sbp",         # Test 1
  "GFP", "SBP",                 # Test 2
  "CGC1", "SCRM1KO",            # Test 3
  "pL4440_gfp", "pL4440_sbp"    # Test 4
))

# Step 3: Compute summary statistics
summary_data <- plot_data %>%
  group_by(Test, Treatment) %>%
  summarise(Mean = mean(Radius), SE = sd(Radius) / sqrt(n()), .groups
= 'drop')

# Step 4: Run t-tests within each Test group
t_test_results <- plot_data %>%
  group_by(Test) %>%
  summarise(
    p_value = t.test(Radius ~ Treatment)$p.value,
    .groups = 'drop'
  ) %>%
  mutate(
    p_label = paste0("p = ", formatC(p_value, format = "f", digits =
3))
  )

# Step 5: Bracket + p-value annotation data
bracket_data <- plot_data %>%
  group_by(Test) %>%
  summarise(y_max = max(Radius), .groups = 'drop') %>%
  left_join(t_test_results, by = "Test") %>%
  mutate(
    y_start = y_max + 0.02 * y_max,
    y_end = y_start + 0.015 * y_max,
```

```
    y_label = y_end + 0.06 * y_max,  # ⬅️ Raised further
    x_start = 1,
    x_end = 2,
    x_label = 1.5
  )

# Step 6: Plot
ggplot(plot_data, aes(x = Treatment, y = Radius, color = Treatment)) +
  geom_jitter(size = 2, alpha = 0.8, width = 0.2, height = 0) +

  geom_point(
    data = summary_data,
    mapping = aes(x = Treatment, y = Mean),
    inherit.aes = FALSE,
    position = position_dodge(width = 0.6),
    shape = 95, size = 5, color = "black"
  ) +

  geom_errorbar(
    data = summary_data,
    mapping = aes(x = Treatment, ymin = Mean - SE, ymax = Mean + SE),
    inherit.aes = FALSE,
    position = position_dodge(width = 0.6),
    width = 0.3,
    color = "black"
  ) +

  # Brackets
  geom_segment(
    data = bracket_data,
    aes(x = x_start, xend = x_end, y = y_end, yend = y_end),
    inherit.aes = FALSE,
    color = "black"
  ) +
  geom_segment(
    data = bracket_data,
    aes(x = x_start, xend = x_start, y = y_start, yend = y_end),
    inherit.aes = FALSE,
    color = "black"
  ) +
  geom_segment(
    data = bracket_data,
    aes(x = x_end, xend = x_end, y = y_start, yend = y_end),
    inherit.aes = FALSE,
    color = "black"
  ) +

  # Rotated p-value text, placed higher
  geom_text(
    data = bracket_data,
```

```
    aes(x = x_label, y = y_label, label = p_label),
    inherit.aes = FALSE,
    size = 4,
    angle = 90
  ) +

  labs(
    x = NULL,
    y = "Radius of Movement Wave Pattern",
    title = ""
  ) +
  theme_minimal() +
  theme(
    panel.grid = element_blank(),
    panel.background = element_rect(fill = "white", color = NA),
    plot.background = element_rect(fill = "white", color = NA),
    axis.line = element_line(color = "black"),
    legend.position = "none",
    axis.text.x = element_text(angle = 90, vjust = 0.5, hjust = 1),
    axis.text.y = element_text(angle = 90, vjust = 0.5, hjust = 0.5),
    strip.text = element_blank()  # ⬅️ Removes Test group facet labels
  ) +
  facet_grid(~Test, scales = "free_x") +
  scale_y_continuous(position = "right") +
  scale_color_manual(values = c(
    "pET_emp" = "#0065CF",
    "pET_sbp" = "#0065CF",
    "GFP" = "#00CAFF",
    "SBP" = "#00CAFF",
    "CGC1" = "#7570b3",
    "SCRM1KO" = "#7570b3",
    "pL4440_gfp" = "#82FFE7",
    "pL4440_sbp" = "#82FFE7"
  ))

# Step 7: Save the plot
ggsave("Radius_Plot_Final.png", width = 10, height = 9, dpi = 300)




##### Ant Code #####
### Aversion Assays ###
library(dplyr)
library(tidyr)
library(ggplot2)
```

```
library(ggsignif)

# Ensure correct factor levels
Aversion_Data$Colony <- factor(
  Aversion_Data$Colony,
  levels = c("Chul5", "Gen17", "LH15", "LH11", "Gen19", "LH14")
)
Aversion_Data$Zone <- factor(Aversion_Data$Zone)
Aversion_Data$Replicate <- factor(Aversion_Data$Replicate)
Aversion_Data$Pair <- factor(Aversion_Data$Pair)
Aversion_Data$Treatment <- factor(
  Aversion_Data$Treatment,
  levels = c("GFP", "PS1", "EMP", "PBS")
)

# Subset data for paired t-tests
gfp_ps1_data <- Aversion_Data %>%
  filter(Treatment %in% c("GFP", "PS1")) %>%
  arrange(Pair, Treatment)

emp_pbs_data <- Aversion_Data %>%
  filter(Treatment %in% c("EMP", "PBS")) %>%
  arrange(Pair, Treatment)

# One-tailed paired t-tests (testing PS1 > GFP and PBS > EMP)
t_test_result_gfp_ps1 <- t.test(
  Adj_z1 ~ Treatment,
  data = gfp_ps1_data,
  paired = TRUE,
  alternative = "less"
)

t_test_result_emp_pbs <- t.test(
  Adj_z1 ~ Treatment,
  data = emp_pbs_data,
  paired = TRUE,
  alternative = "less"
)

# Extract p-values
p_value_gfp_ps1 <- t_test_result_gfp_ps1$p.value
p_value_emp_pbs <- t_test_result_emp_pbs$p.value

# Summary statistics
summary_data <- Aversion_Data %>%
  group_by(Zone, Treatment) %>%
  summarise(
    Mean = mean(Adj_z1),
    SE = sd(Adj_z1) / sqrt(n()),
    .groups = "drop"
```

```
  )

# Y-axis limit and annotation positions
ymax <- 0.36
pval_y_pos_1 <- ymax * 0.8

# Jitter settings
jitter_width <- 0.2
treat_levels <- levels(Aversion_Data$Treatment)
treat_map <- setNames(seq_along(treat_levels), treat_levels)

# Jittered x positions
jittered_data <- Aversion_Data %>%
  mutate(
    Treatment_num = treat_map[as.character(Treatment)],
    x_jitter = Treatment_num + runif(n(), -jitter_width, jitter_width)
  )

# Paired line segment data
line_data <- jittered_data %>%
  group_by(Pair) %>%
  filter(n() == 2) %>%
  arrange(Treatment_num) %>%
  summarise(
    x = x_jitter[1], xend = x_jitter[2],
    y = Adj_z1[1], yend = Adj_z1[2],
    Colony = Colony[1],
    Zone = Zone[1],
    .groups = "drop"
  )

# Final plot
ggplot(jittered_data, aes(x = Treatment, y = Adj_z1, color = Colony,
shape = Colony)) +
  geom_signif(
    comparisons = list(c("GFP", "PS1")),
    annotations = paste0("p = ", signif(p_value_gfp_ps1, 3)),
    textsize = 4,
    y_position = pval_y_pos_1 - 0.3,
    color = "black",
    tip_length = -0.02,
    vjust = 2.5
  ) +
  geom_signif(
    comparisons = list(c("EMP", "PBS")),
    annotations = paste0("p = ", signif(p_value_emp_pbs, 3)),
    textsize = 4,
    y_position = pval_y_pos_1 - 0.3,
    color = "black",
    tip_length = -0.02,
```

```
  vjust = 2.5
) +
geom_segment(
  data = line_data,
  aes(x = x, xend = xend, y = y, yend = yend, color = Colony),
  inherit.aes = FALSE,
  linetype = 1,
  size = 0.5,
  alpha = 1
) +
geom_point(
  data = jittered_data,
  aes(x = x_jitter, y = Adj_z1, color = Colony, shape = Colony),
  size = 2,
  alpha = 1
) +
geom_point(
  data = summary_data,
  aes(x = Treatment, y = Mean),
  inherit.aes = FALSE,
  shape = 95,
  size = 5,
  color = "black"
) +
geom_errorbar(
  data = summary_data,
  aes(x = Treatment, ymin = Mean - SE, ymax = Mean + SE),
  inherit.aes = FALSE,
  width = 0.2,
  color = "black"
) +
facet_wrap(~Zone) +
labs(x = NULL, y = "Ants in the aversion zone (%)") +
theme_minimal() +
theme(
  legend.position = "none",  # <-- this removes the legend
  panel.grid = element_blank(),
  panel.background = element_rect(fill = "white", color = NA),
  plot.background = element_rect(fill = "white", color = NA),
  axis.line = element_line(color = "black"),
  axis.text.x = element_text(angle = 0, hjust = 0.5),
  axis.text.y = element_text(angle = 0)
) +
scale_color_manual(values = c(
  "Chul5" = "#FF0000",
  "Gen17" = "#B50406",
  "LH15" = "#FFC4E0",
  "LH11" = "#E47B7B",
  "Gen19" = "#C25200",
  "LH14" = "#65102A"
```

```
  )) +
  scale_shape_manual(values = c(
    "Chul5" = 16,
    "Gen17" = 17,
    "LH15" = 15,
    "LH11" = 18,
    "Gen19" = 8,
    "LH14" = 4
  )) +
  coord_cartesian(ylim = c(-0.05, ymax))

# Save plot as image
ggsave("aversion_plot_merged.png", width = 4, height = 6, dpi = 1000)
```

```
### Aversion Assay Time Course ###
# Load necessary libraries
library(dplyr)
library(ggplot2)

# Summarize the data: calculate mean for Ants
Timecourse_Summary <- Timecourse_Data %>%
  group_by(Colony, Treatment, Timepoint, Replicate) %>%  # Include
Replicate in grouping
  summarise(
    mean_Ants = mean(Ants, na.rm = TRUE),
    .groups = 'drop'
  ) %>%
  filter(
    is.finite(mean_Ants),
    !is.na(Timepoint)
  )

# Plot: 4 lines for EMP, GFP, PS1, SBP — styled by Treatment
ggplot(Timecourse_Summary, aes(x = Timepoint, y = mean_Ants)) +
  geom_point(aes(color = Treatment), size = 1.5, alpha = 0.01) +
  geom_smooth(aes(color = Treatment, linetype = Treatment), method =
"loess", se = FALSE, size = 1) +
  scale_color_manual(values = c(
    "GFP" = "#FF0000",
    "PS1" = "#FF0000",
    "EMP" = "#B22222",
```

```
    "SBP" = "#B22222"
  )) +
  scale_linetype_manual(values = c(
    "GFP" = "solid",
    "PS1" = "dotdash",
    "EMP" = "solid",
    "SBP" = "dotdash"
  )) +
  labs(
    title = "Timecourse",
    x = "Time (s)",
    y = "Ants within the aversion zone (%)",
    color = "Treatment",
    linetype = "Treatment"
  ) +
  theme_minimal() +
  theme(
    panel.grid.major = element_blank(),
    panel.grid.minor = element_blank(),
    text = element_text(size = 14),
    legend.position = "right",
    legend.text = element_text(size = 14),
    legend.title = element_text(size = 16, face = "bold"),
    legend.key.size = unit(1.5, "lines")
  ) +
  coord_cartesian(ylim = c(0, 0.4))  # Set y-axis limit without
removing data

# Save plot as image
ggsave("aversion_timeplots_merged.png", width = 12, height = 6, dpi =
300)
```

```
### Antennation Assay ###
# Load required library
library(ggplot2)

# Filter only CC and TT treatments
filtered_data <- subset(Antennation, Treatment %in% c("CC", "TT"))

# Create violin plot WITH custom shapes and bigger points
ggplot(filtered_data, aes(x = Treatment, y = Count, fill = Treatment))
```

```
+
  geom_violin(trim = FALSE, color = "black") +
  geom_boxplot(width = 0.1, outlier.shape = NA) +
  geom_jitter(
    aes(color = Colony, shape = Colony),
    width = 0.15,
    alpha = 0.7,
    size = 4   # Points twice as big
  ) +
  theme_minimal(base_size = 14) +
  theme(
    legend.position = "none",
    plot.title = element_blank(),
    plot.subtitle = element_blank(),
    panel.grid.major = element_blank(),
    panel.grid.minor = element_blank()
  ) +
  labs(
    x = "Treatment",
    y = "Count"
  ) +
  scale_fill_manual(
    values = c("CC" = "#FFFFFF", "TT" = "#FFC20A")
  ) +
  scale_color_manual(
    values = c(
      "LH14" = "#6F0029",
      "LH11" = "#E47B7B",
      "Chul5" = "#FF0000"
    )
  ) +
  scale_shape_manual(
    values = c(
      "Chul5" = 16,  # Solid circle
      "LH11" = 18,   # Solid diamond
      "LH14" = 4     # Cross
    )
  )

# Save plot as image
ggsave("antennation_violin.png", width = 9, height = 6, dpi = 1000)
```

```
### Nest Behavior Statistical Analysis ###
### Statistical Analysis ###
# Load libraries
library(lme4)
library(lmerTest)
library(ggplot2)

# Use your loaded dataset
data <- Nest_Behavior_Master_Sheet

# Convert to factors
data$Treatment <- as.factor(data$Treatment)  # 'C' or 'T'
data$Colony <- as.factor(data$Colony)

# Assign Individual ID — 20 individuals × 600 rows
data$Individual <- as.factor(rep(1:20, each = 600))

# Fit linear mixed-effects model with NNDistance as response
model <- lmer(NNDistance ~ Treatment + FRAME + (1 | Colony/
Individual), data = data)

# View model summary
summary(model)

# Plot mean NNDistance over time by treatment
ggplot(data, aes(x = FRAME, y = NNDistance, color = Treatment)) +
  stat_summary(fun = mean, geom = "line", size = 1) +
  labs(title = "Average NNDistance Over Time by Treatment",
       x = "Frame (Time)", y = "NNDistance from Start") +
  theme_minimal()

# Fit the original model without interaction
model_no_interaction <- lmer(NNDistance ~ Treatment + FRAME + (1 |
Colony/Individual), data = data)

# Fit the model with Treatment × FRAME interaction
model_interaction <- lmer(NNDistance ~ Treatment * FRAME + (1 |
Colony/Individual), data = data)

# View the model summary for the interaction model
summary(model_interaction)

# Compare the two models using a likelihood ratio test
anova(model_no_interaction, model_interaction)

# Plot average NNDistance over time by Treatment group
ggplot(data, aes(x = FRAME, y = NNDistance, color = Treatment)) +
  stat_summary(fun = mean, geom = "line", size = 1) +
  labs(title = "NNDistance Over Time by Treatment",
```

```
      x = "Frame (Time)", y = "NNDistance") +
  theme_minimal()
```

```
### Difference Area Plot ###
library(dplyr)
library(ggplot2)
library(tidyr)
library(zoo)

# Step 1: Summarize and smooth
df_summary <- Nest_Behavior_Master_Sheet %>%
  group_by(FRAME, Treatment) %>%
  summarise(mean_dist = mean(NNDistance / 7.35, na.rm = TRUE), .groups
= "drop") %>%
  pivot_wider(names_from = Treatment, values_from = mean_dist)

df_diff <- df_summary %>%
  rename(T_val = T, C_val = C) %>%
  mutate(
    diff = T_val - C_val,
    diff_smooth = rollmean(diff, k = 5, fill = NA, align = "center"),
    FRAME_s = FRAME / 2
  ) %>%
  filter(!is.na(diff_smooth)) %>%
  arrange(FRAME_s)

# Step 2: Tag positive/negative regions
df_diff <- df_diff %>%
  mutate(sign = ifelse(diff_smooth >= 0, "pos", "neg"))

# Step 3: Group consecutive same-sign values using data.table rleid
df_diff <- df_diff %>%
  mutate(segment = data.table::rleid(sign))

# Step 4: Build ggplot with one ribbon per segment
ggplot() +
  geom_hline(yintercept = 0, linetype = "dashed", color = "gray") +

  # Positive ribbons (above 0)
  geom_ribbon(
    data = df_diff %>% filter(sign == "pos"),
```

```
    aes(x = FRAME_s, ymin = 0, ymax = diff_smooth, group = segment),
    fill = "#65102A", alpha = 1.0
  ) +

  # Negative ribbons (below 0)
  geom_ribbon(
    data = df_diff %>% filter(sign == "neg"),
    aes(x = FRAME_s, ymin = diff_smooth, ymax = 0, group = segment),
    fill = "#FF0000", alpha = 1.0
  ) +

  # Smoothed line on top
  geom_line(
    data = df_diff,
    aes(x = FRAME_s, y = diff_smooth),
    color = "#000000", linewidth = 0.5
  ) +

  scale_x_continuous(
    limits = c(0, 300),
    breaks = seq(0, 300, by = 30)
  ) +
  labs(
    x = "Time [s]",
    y = "Smoothed Distance Difference [mm]"
  ) +
  theme_minimal() +
  theme(
    panel.grid.major = element_blank(),
    panel.grid.minor = element_blank()
  )

# Step 5: Save the plot
ggsave("Nearest_Neighbor_Plot.png", width = 10, height = 5, dpi = 300)
```
