## Supplementary material for "The First of Us: *Ophiocordyceps* use a novel scramblase-binding peptide to manipulate zombie ants": Data S9

Codon optimization of Ophcf2|06345 for *C. elegans*

**Transcript (SP region)**

**ATGAACAAGCTCTTCCTCGTCATCATCTCCCTGGTGTGCGCCGCCGCCGCTCTCCCCGCGCAGCAGGCCAACGCC**ATCGACGCCAGCCGATCCTTTACCTGTCCGTCGCGCGTGCTGGCGTTTTGCCGCGCGAGGAACGTCCATTCCGGCTGTACCTTTAACGGTAGCTTCCGGTCTGATGCTATGGAGACCTGTAGGGAGTGTCACTGTGGG**TGA**

**Output (spp-1 SP)(no synthetic intron)**

**ATGACTCGCATTCTTCCGTGTCTTTTCCTTGTCTTGCTCGCCGCTGCTCCACTTCTAGCC**ATCGATGCTTCTCGTTCTTTCACCTGCCCATCTCGTGTTCTTGCTTTCTGCCGTGCTCGTAACGTTCACTCTGGATGCACTTTCAACGGATCTTTCCGTTCCGACGCCATGGAGACTTGCCGCGAATGCCACTGCGGA**TAA**

Note: Sequence is too short for a synthetic intron

**Optimization Details**

ORIGINAL    Codon Adaptation Index: 0.20   GC content: 59%  
OPTIMIZED   Codon Adaptation Index: 0.86   GC content: 52%  

ORIGINAL    ATC GAC GCC AGC CGA TCC TTT ACC TGT CCG TCG CGC GTG CTG GCG TTT TGC CGC GCG AGG AAC GTC CAT TCC GGC TGT ACC TTT AAC GGT 
OPTIMIZED   ATC GAT GCT TCT CGT TCT TTC ACC TGC CCA TCT CGT GTT CTT GCT TTC TGC CGT GCT CGT AAC GTT CAC TCT GGA TGC ACT TTC AAC GGA 
AMINO ACIDS Ile Asp Ala Ser Arg Ser Phe Thr Cys Pro Ser Arg Val Leu Ala Phe Cys Arg Ala Arg Asn Val His Ser Gly Cys Thr Phe Asn Gly 

ORIGINAL    AGC TTC CGG TCT GAT GCT ATG GAG ACC TGT AGG GAG TGT CAC TGT GGG TGA 
OPTIMIZED   TCT TTC CGT TCC GAC GCC ATG GAG ACT TGC CGC GAA TGC CAC TGC GGA TAA 
AMINO ACIDS Ser Phe Arg Ser Asp Ala Met Glu Thr Cys Arg Glu Cys His Cys Gly Ter

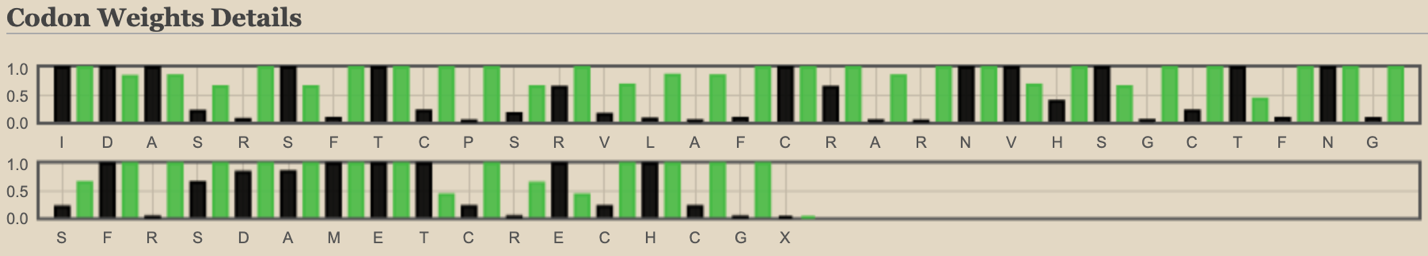
